# DIANNE: Segmentation-Free Localization of Histology Differential Attributes

**DOI:** 10.64898/2026.04.28.721103

**Authors:** Sergii Domanskyi, Jill C. Rubinstein, Todd B. Sheridan, Adam Thiesen, Javad Noorbakhsh, Juliana Alcoforado Diniz, Ramalakshmi Ramasamy, Dylan S. Baker, Riley Sheldon, Qian Wu, George Kuchel, Nicolas Musi, Sara E. Espinoza, Francisco G. Cigarroa, Paul Robson, Jeffrey H. Chuang

## Abstract

Pathologist-guided distinctions within histology and spatial omic images provide insights into health and disease, with digital pathology leveraging artificial intelligence to automate such assessments. To train computational models, current digital pathology methods rely on upfront manual annotations, which are time-consuming to generate. Pre-annotation is poorly suited to investigating novel spatial behaviors—a major need driven by advances in spatial profiling—for which annotation criteria and data needs will be uncertain. To address these challenges, we present DIANNE, a digital pathology annotation support and discovery approach for rapid training and inference of spatial differential attributes, enabled by train-time Positive Class Mixup Augmentation. DIANNE can compute foundation model-derived segmentation-free localization of differential classifiers across whole slide H&E images within seconds on a workstation, enabling interactive investigation of spatial niches. Predictive models can be re-trained in real-time in response to patch or regional annotation changes, clarifying determinative biological attributes across slides from only a few dozen annotated patches. We demonstrate the effectiveness of DIANNE for tumor detection, artifact identification, and exploration of pancreatic, fetal membranes and kidney tissue structures. DIANNE also provides analogous capabilities for IHC, multiplex immunofluorescence, and registered spatial transcriptomic+H&E images. DIANNE is implemented in a Jupyter toolkit, enabling rapid development of high-resolution classifiers from weakly-supervised training. DIANNE provides a practical system to quantitatively understand known and novel spatial phenotypes.

## Introduction

Understanding the differences among and within histology images is crucial for clinical pathology and biomedical scientific discovery [1,2,3]. Pathologists assess these differences to make diagnoses, determine the severity of diseases, and guide treatment plans to improve patient outcomes [4]. Scientific insight is gained through the process of defining, validating, and refining classifiers of known and newly considered histological states—such as those observed in cancer, inflammation, infections, or genetic disorders. Digital pathology enhances these activities through automated analysis of digitized slides, notably through deep learning (DL) and other artificial intelligence (AI) and machine learning (ML) methods for assessment of tissue morphology, biomarker detection, and disease classification [5,6].

Multiple sources of technical complexity can hinder the learning of biologically important structures from whole slide images (WSI), such as hematoxylin and eosin (H&E), multiplex immunofluorescence (mIF), or immunohistochemistry (IHC) images. For example, different microscope scanners have varying resolutions and color profiles, and images are affected by protocols for sample collection, preservation, tissue sectioning, staining, and imaging conditions, leading to batch effects and data inconsistency [7,8]. Image harmonization can mitigate technical biases, but different histological structures may vary in spatial complexity, making it challenging to develop a slide-level harmonization optimal for all classification tasks. Flexibility and efficiency in dealing with such challenges are therefore crucial, especially when investigating previously unknown spatial structures.

Here we address such challenges by developing a data augmentation approach that is fast enough to enable real-time interactive classifier optimization for flexible and accurate prediction of biological structures in tissue images. Our approach is inspired by the “selective mixup” technique [9,10], in which new training samples and labels are generated as a combination of existing samples. We perform this mixup in the SAMPLER data representation, which represents each image via the cumulative distribution functions of image-tile features, providing important advantages [11]. Specifically, we develop train-time Positive Class Mixup Augmentation (PCMA), in which we generate augmented samples by combining SAMPLER representations of images from positive and negative classes. This augmentation yields improved feature selection in a latent space known to enable efficient, accurate classification.

To share these capabilities, we have developed the software package DIANNE: <u>D</u>ifferential Image <u>Ann</u>otator <u>E</u>nvironment. DIANNE is implemented in a Jupyter-based toolkit and contains four workflows developed with PCMA. The four workflows stem from two choices: static/interactive and histology/molecular. The static/interactive decision specifies whether classifier training is based on a fixed set of annotated images, or whether the user interactively trains classifiers. The histology/molecular decision specifies whether the images are conventional histological image types (H&E, mIF, or IHC), or are molecular spatial transcriptomic data registered to histology images. DIANNE implements current image processing pipelines [12] as well as novel Multiplex Immunofluorescence Extraction (MIE) data processing pipelines to handle these image types, see Methods. These four workflows enable rapid training of classifiers to generate spatially resolved differential attribute probability heatmaps.

The speed and accuracy of DIANNE enhance hypothesis-free, exploratory analysis, allowing researchers to interactively develop classifiers from tissue image datasets without requiring uncertain and laborious pre-annotation. This exploratory capability is important for spatial studies, whose scope can be challenging to specify, especially for previously unknown physiological mechanisms. By bridging complex imaging data and model training with intuitive visual annotation, DIANNE empowers users to rapidly uncover novel spatial biology and clinically predictive spatial features.

## Results

### DIANNE Overview: Multi-Modal Workflows for Image Attribute Localization

Effective training of image classifiers is constrained by properties of the data representation and the training data. DIANNE provides improved image classification by integrating the recently developed quantile-based SAMPLER representation with a distinctive data augmentation procedure. Given an input set of images with class labels, DIANNE can efficiently augment them and train a classifier on the augmented data.

The first step in DIANNE is to embed each image (either a WSI or a multi-tile patch) into the SAMPLER representation. To do this, the image is decomposed into fixed-size tiles that are input into a pre-specified foundation model, e.g. [13–16], yielding deep learning features for each tile. The SAMPLER representation is then computed from the quantile distributions of the deep learning features across all tiles in the image.

The second step in DIANNE is to augment the training data. Class mixup data augmentation techniques have been developed in other contexts to improve training by mixing different classes, providing improved feature selection for distinction of positive and negative classes [9,10]. The effectiveness of class mixup depends on whether the augmentation preserves biological behaviors informative for classification. For instance, tumor-positive histology slides typically contain some normal tissue regions in contiguous blocks on the margins around the tumor. Such spatial tendencies can be informative, so we have developed an augmentation approach that incorporates distributional relationships among proximate tiles. Naive augmentation based on independent tile sampling would ignore such relationships, producing augmented datasets that superpose tumor and normal signal in a semantically inconsistent manner. DIANNE improves upon this by performing class mixup in the SAMPLER latent space, whose distribution-based formulation provides superior semantic coherence. Importantly, the SAMPLER representation supports mixup computation without the need to re-process individual tiles, greatly increasing computational efficiency. Details are available in the Methods.

DIANNE provides four workflows, which we summarize here and detail below. All workflows begin with standardized processing of WSIs. The STQ [12] pipeline normalizes H&E and IHC slides, generates tile grids, and extracts imaging features via histopathology foundation models. The MIE pipeline processes mIF WSIs. Imaging features are extracted using pre-trained histopathology foundation models, including CTransPath [13], MoCoV3 [14], UNI/UNI2 [15], and CONCH [16] for H&E data, and KRONOS [17] for mIF spatial proteomics data, though alternative models can be swapped in (see Methods).

The *<u>histology static workflow</u>* applies the SAMPLER/PCMA approach to a set of histology images (H&E, IHC, or mIF) with pre-specified (“static”) annotations, a traditional input for current digital pathology studies. This workflow is designed for weakly supervised training from image-level annotations, e.g., “cancer tissue slide”, “normal tissue slide”, “tissue slide with tissue folds artifact” or “slide clear of any artifacts”, which are more commonly available than pixel-level annotations (**Fig. 1c**).

**Figure 1.**
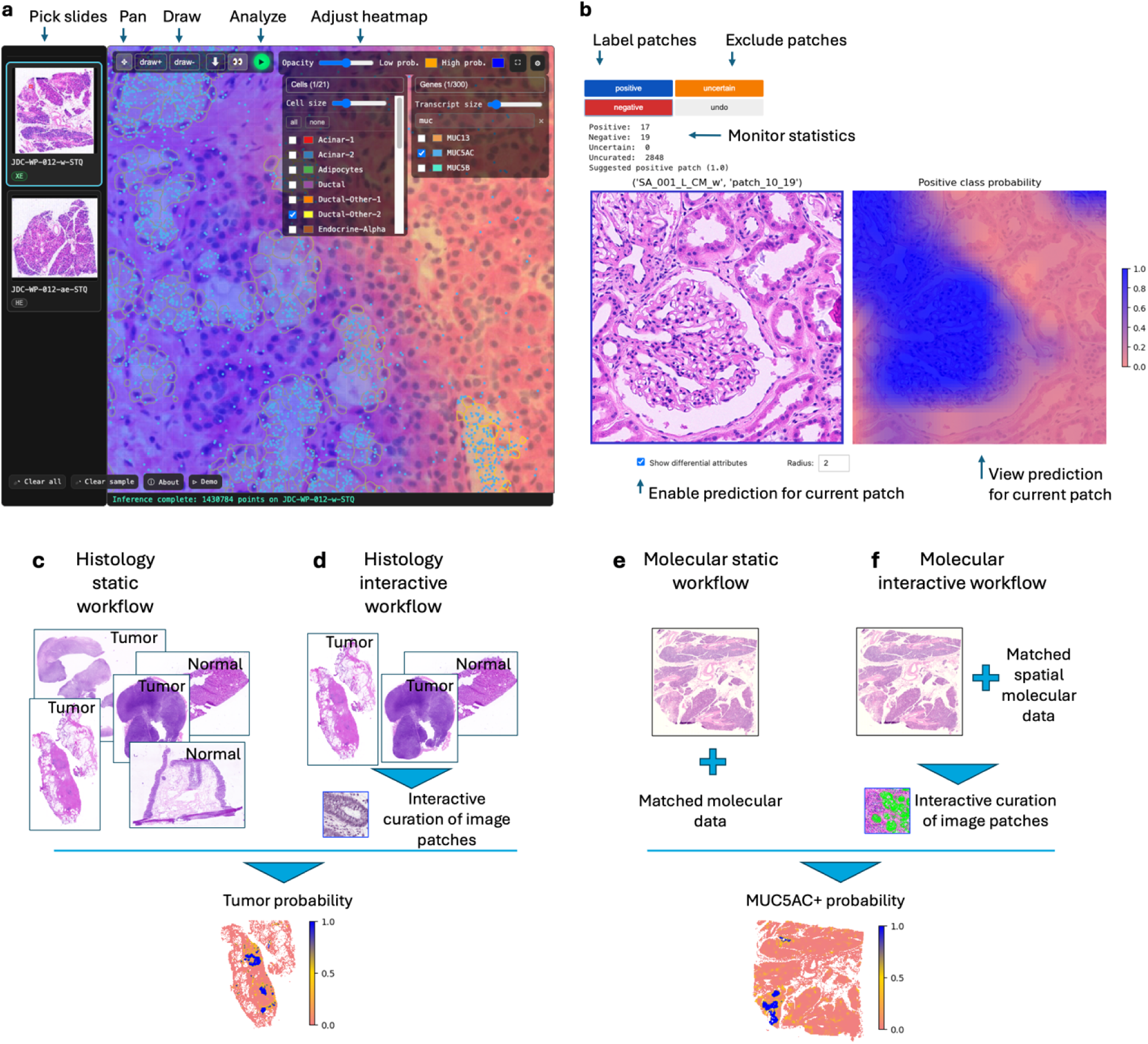
DIANNE graphical user interface (GUI) and workflows. **a**, Freehand GUI allows the user to navigate to a single whole slide image by clicking on the thumbnail on the left and then by zooming and panning the user draws positive or negative contours to label regions of interest based on their own visual judgment. The blue-orange semi-opaque heatmap shows the DIANNE predictions of pancreatic intraepithelial neoplasia (PanIN). When available, as indicated on the thumbnails, the spatial transcriptomics data is overlaid on the H&E-stained slide image, here transcripts MUC5AC are shown in blue, and cell boundaries are overlaid over the image. **b**, Guided GUI presents image patches for user curation to label the patches as positive or negative, here an H&E image patch of human kidney with apparent glomeruli (left panel), with the model highlighting regions of high glomerular tissue probability (right panel). **c**, Histology static histology workflow. SAMPLER representations and class labels are used for classifier training. The classifier is then used for spatial inference, e.g., tumor/normal prediction. The static workflow can operate either on whole slides or on patches of tiles. **d**, Histology interactive workflow. Interactive annotation enables the operator to retrain classifiers and display predicted label status of patches in real time. Interactive workflows operate on patches of tiles. **e**, Molecular static workflow. This workflow builds classifiers from histology images aligned to spatial molecular data (e.g. spatial transcriptomics). It can also be applied to sample-matched non-spatial molecular data. **f**, Molecular interactive workflow. This workflow provides interactive spatial molecular classifier training. It facilitates this process by displaying molecular data on the histology image patches during interactive annotation. Here MUC5AC Xenium transcripts (green) are overlaid on a human pancreas H&E. All four workflows process WSIs with the STQ pipeline (or MIE pipeline for mIF images) to normalize images, make a grid of tiles, and extract imaging features for each tile. These features are used to generate a SAMPLER representation followed by PCMA data augmentation and training of a classifier that can output spatially resolved differential attribute probability heatmaps.

The *<u>histology interactive</u>* workflow uses the SAMPLER/PCMA approach for real-time labeling and re-training of spatial classifiers on WSIs (**Fig. 1d**). It operates on patches of tiles (default 8×8). The interactive toolkit, **Fig. 1a-b**, allows examination and designation of WSI image patches as positive or negative. The next patch to be annotated is selected based on the current classifier via an active learning algorithm, enabling rapid human-in-the-loop curation, **Supplementary Fig. 1**. A demonstration of the histology interactive workflow is provided in **Video S1 and S2**.

The *<u>molecular static</u>* workflow applies to spatial transcriptomics data (e.g. Visium, Xenium, Atera) with matched histology H&E-stained images. This workflow trains a classifier that identifies associations between spatial molecular features (e.g. genes, pathways, or cell types) and H&E image features [18,19,20]. The workflow derives patch labels from the molecular data (**Fig. 1e**). It can also operate on molecular data derived from non-spatially resolved assays.

The *<u>molecular interactive</u>* workflow enables real-time labeling and re-training of classifiers connecting spatial molecular features with histology images. It assists the operator in making patch annotations by overlaying the molecular data on histology image patches (**Fig. 1f**). This functionality enables users to easily incorporate both molecular markers and morphology into classifier training for newly identified multiplex tissue structures. A demonstration of the molecular interactive workflow is provided in Video S3 and S4.

### Evaluation of Histology Static Workflow Tumor / Normal Detection in Pediatric Sarcoma

To assess the histology static workflow, we trained and evaluated classifiers on a tumor/normal set. Tumor samples comprise 887 pediatric sarcoma images [21]. The normals are 310 normal TCGA [22] images from specimens of solid tissues (sample type code 11A), as well as 31 normal lymph node images from CAMELYON16 [23] included to distinguish non-tumor lymphoid aggregates. We used DIANNE to process WSIs through the STQ pipeline, which creates a grid of tiles and extracts CTransPath [13] foundation model imaging features for each tile. Slide-level SAMPLER representations were used as inputs for logistic classifier training, with each slide designated as either positive or negative. PCMA was used to augment the training data in the SAMPLER latent space.

To evaluate DIANNE, we trained a tumor/normal classifier on 857 of the sarcoma and 291 of the normal whole slide images using slide-level labels (15 seconds, 1 CPU). A clinical pathologist annotated the remaining 30 sarcoma slides and 50 normal slides with tile-resolution tumor fraction, providing a validation set. **Fig. 2a** shows the pathologist annotations for case RMS2234 [21]. The corresponding DIANNE tumor probability heatmap is shown in **Fig. 2b**. We inspected the normal slides and found that a few (2 of 50 validation slides; 7 of 291 training slides) contained predicted tumor hotspots, i.e. false positives. However, most such false positives were due to local tissue distortions or imaging artifacts, as confirmed by the clinical pathologist. We then compared DIANNE to other recent digital pathology approaches. The CLAM pipeline [24] (**Fig. 2c**) uses a contrastive learning approach without augmentation. It yielded lower specificity and higher recall when trained on the same dataset and the same STQ-extracted features (30 minutes, A100 GPU). We also compared to Segmenter [25,26] (**Fig. 2d**), a pixel-level ViT-based segmentation model. Because Segmenter requires pixel-level annotations for training, we annotated an additional 70 tumor slides and 14 normal slides and trained via fine-tuning on these. Segmenter yielded higher resolution spatial predictions but required more computational resources and training time (6 hours, A100 GPU). These results are summarized in **Fig. 2e**.

**Figure 2.**
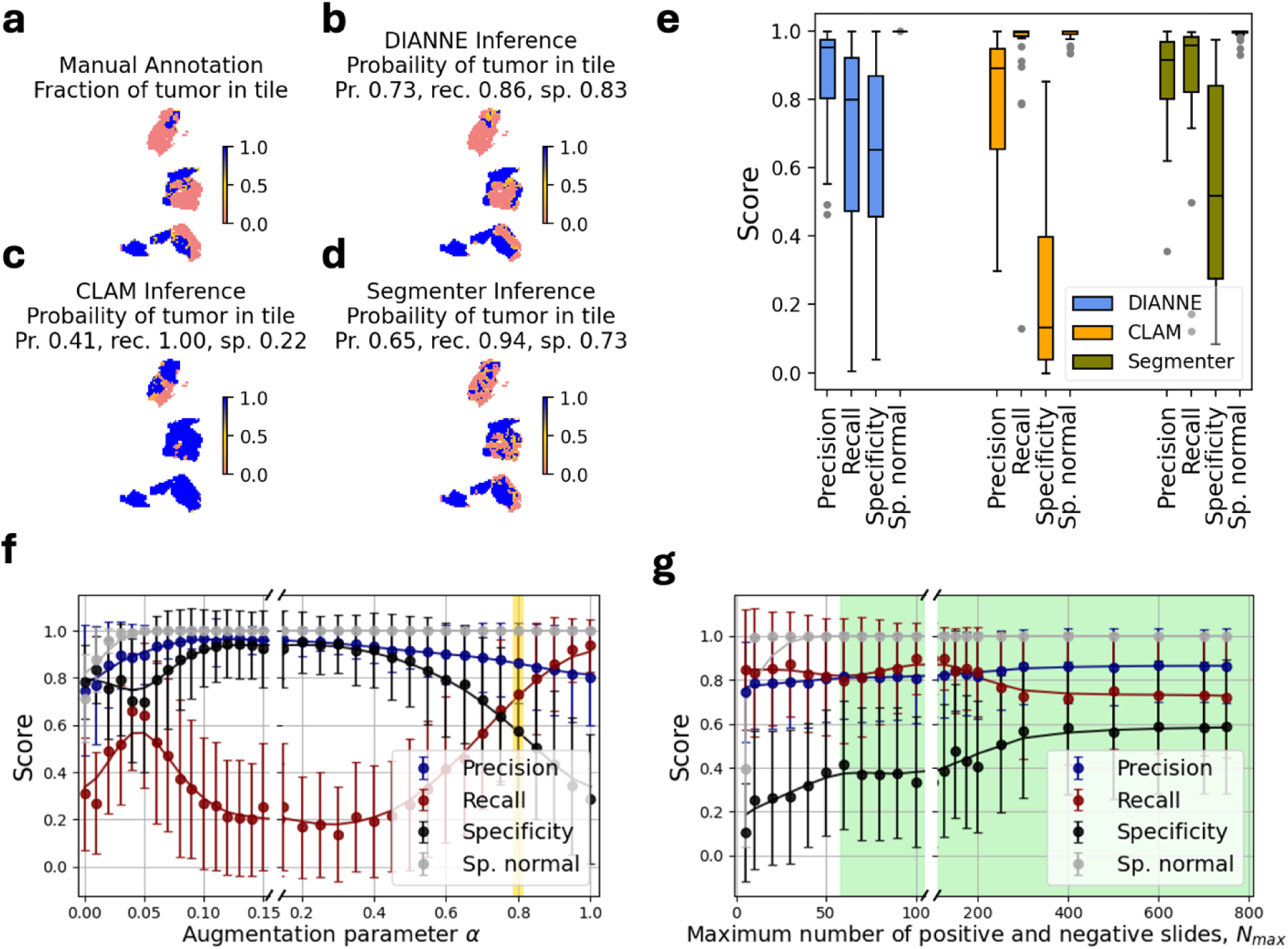
Spatial inference of tumor/normal for pediatric sarcomas. **a**, Pathologist annotation of tumor fraction in case RMS2234 [21]. **b-d**, DIANNE-, CLAM- and Segmenter-inferred heatmaps of predicted regional tumor probability. **e,** Classification metrics for DIANNE, CLAM, and Segmenter. “Specificity” refers to specificity-on-tumor-containing-slides, and “Sp. normal” refers to specificity-on-normal-slides. DIANNE training was done with augmentation parameter =0.8, the optimal value determined from plot **f. f,** DIANNE classification metrics as a function of augmentation parameter. The optimal parameter value is highlighted in yellow. =1 corresponds to no augmentation**. g**, DIANNE classification metrics as a function of training size. Error bars are derived from replicate sampling.

As an alternative to SAMPLER, we tested a moment-based representation using the first 10 raw moments, *{(x^k^_i_)_i_}^10^_K=1_*, of tile deep learning features, *x*. The moment-based classifier yielded recall of 1.000 on the validation slide but at the expense of low specificity, regardless of the augmentation parameter value. The medians of precision/specificity-on-tumor-containing-slides/specificity-on-normal-slides across the validation set were 0.852/0.028/0.999 for a=1 (no augmentation), 0.843/0.023/0.996 for a=0.8, 0.834/0.008/0.977 for a=0.5, and 0.833/0.003/0.866 for a=0.2. We observed that the means of tile imaging features were the dominant predictive features. Due to the inferior performance of the moment-based approach, we used SAMPLER for all subsequent tasks.

We next tested whether aggregating multiple normal slides into each SAMPLER normal representation could increase performance α. We hypothesized that combining slides could improve predictions by emphasizing shared features; however, this approach could also suffer from slide-specific batch effects. To construct aggregated slide representations, we randomly selected S normal slides and sampled 2,000 tiles from these to generate a normal SAMPLER representation. This procedure for generating a normal representation was used to augment tumor samples during model training. When S = 1, the approach corresponds to the baseline configuration previously employed, **Supplementary Fig. 2a**. Performance was evaluated for S=2, 3, 4, 5, 10, 15, and 30 on the validation set. For S = 2 and 3, classification performance remained comparable to the baseline. However, when S = 4 or 5, the specificity began to degrade, and S = 15 resulted in a zero specificity. Thus, when tiles are sampled from many slides, the SAMPLER representation becomes dominated by slide-specific batch effects, reducing its utility for augmentation.

Patch level aggregation analysis reinforced these findings. Using pathologist annotated sarcoma slide RMS2341, we subdivided the tissue into patches, **Supplementary Fig. 2b**. We then generated aggregated representations by combining P sampled negative patches from the slide. Increasing P beyond 1 produced a notable decline in performance (**Supplementary Fig. 2c**). Thus, the effectiveness of PCMA is dependent on using individual negative samples to augment individual positive samples. This is because the distributional properties of an aggregated set of patches are not equivalent to the distributional properties within individual patches, which reflect local behaviors.

We also investigated DIANNE’s performance as a function of the augmentation parameter. We found that = 0.8 yielded optimal performance with median precision 0.953, recall 0.800, specificity on tumor slides 0.652 and specificity on normal slides 1.0 (**Fig. 2f**). Note that the moderate specificity on tumor slides is due to the fact that tumor slides usually contain significant amounts of normal tissue. Without augmentation (α = 1) there was lower precision 0.897, high recall 0.991 and greatly degraded specificity (on tumor slides) 0.176, though median specificity remained at 1.0 on normal slides. Notably, DIANNE achieved nearly optimal performance with even small training sets. For example, it obtained precision 0.819, recall 0.797 on tumor slides 0.433 when trained on 60 randomly selected slides from the full training set (**Fig. 2g**). Thus the histology static workflow provides fast, accurate training of classifiers with modest training set needs.

### Analysis of Pancreas Using the Histology Static Workflow

We used DIANNE to analyze a system where spatial structures are not well-known and classification tasks are uncertain, a common situation in spatial omic analysis. In particular, we considered a set of 32 pancreas slides. Gross anatomical regions (head, tail) are commonly used to organize pancreas tissue analysis [27,28], as local region-specific structures are not fully understood. We used DIANNE to train a patch-level classifier of head/tail status based on H&E attributes. Each patch consisted of an 8×8 grid of tiles, with each tile 224×224 pixels. The classifier was trained from the patch SAMPLER representations computed from the tiles within each patch. We achieved 58% accuracy and 0.50 F1 score with leave-one-donor-out cross-validation, indicating poor discrimination of head/tail status given the class prevalence (head: tail = 0.54: 0.46).

The poor accuracy of the head/tail classifier appears to be due to the limitations of gross category definitions that ignore intra-region heterogeneity. For example, we observed a number of “head-like” features—such as ductal density and acinar organization— in the tail region of the pancreas, **Supplementary Fig. 3**. Similarly, we analyzed classifiers trained on age. We trained classifiers using donor age group (young and old) as patch labels (5 young, under 33 years; 4 old, above 50 years. 2-3 tissue images per donor; total 20 images) and PCMA augmentation parameter 0.65. Leave-one-donor-out cross-validation yielded 65% accuracy and 0.63 F1 score.

This performance exceeds the 50% expected from class prevalence, indicating that age associated features are present. However, young donors also contained morphologies associated with older pancreases (0.1-0.3% of the tissue tiles, **Supplementary Fig. 3 f-h**). These findings show the need for improved tools to discover region-specific histological patterns.

### Exploratory Analysis with the Histology Interactive Workflow

To address the need for improved discovery tools, we developed the interactive histology workflow with a guided graphical user interface, **Fig. 1b**. This workflow provides speed, scalability, and interactivity to enable efficient exploratory investigations. After initialization, the tool loads preprocessed imaging features (∼1 second per 1 cm^2^ tissue at 0.25 mm per pixel) and generates patch representations in the same time scale. The viewport can load and display image patches in < 1 second. This efficiency supports interactive classifier training important for discovery analysis, in which training data needs will not be *a priori* known. Users curate patches iteratively, with updated predictions displayed immediately for the next patch. Whole slide inferences can also be computed and displayed on demand (compute time: <60 seconds per slide, or 5 seconds per 1 cm^2^ tissue at 0.25 mm per pixel on a 4-core CPU). The DIANNE Jump-start tool, **Supplementary Fig. 4**, assists in the initiation of training by summarizing the morphological diversity in the dataset. It displays a representative patch from each of N clusters, derived via KMeans clustering using patch correlation distances. The operator selects a patch to seed the annotation tool and begin interactive classifier training. Examples demonstrating this workflow are described below, with validations on slides annotated by a clinical pathologist (see Methods).

### Interactive classification of pancreatic intraepithelial neoplasia

We first trained a classifier for pancreatic islets, which are critical to endocrine function and diabetes pathology [31,32]. To do this, we interactively curated positive and negative patches from a histological section from a pancreas tail tissue block from a 23-year-old female donor (total tissue size: 987 patches). **Fig. 3** shows classifier predictions based on interactive user labeling of 41 positive and 41 negative patches. The classifier successfully detected islets apparent in the H&E (**Fig. 3a**), often with probability values >0.9 (**Fig. 3b**). Predicted islets were dispersed over the complete tissue. We validated the classifier based on pixel-level manual pathologist annotations for the whole slide. This yielded AUROC 0.937. The AUPRC was 0.438, considerably better than random expectations given class imbalance, as pathologist-verified islets cover only 1.5% of pixels in the slide. We repeated interactive training using a pancreas head tissue block from the same donor, generating a classifier by annotating 13 positive and 20 negative patches (out of 1337 total patches). The resulting classifier also yielded strong validations on the pathologist annotations for the tail block, with AUROC 0.927 and AUPRC 0.403.

**Figure 3.**
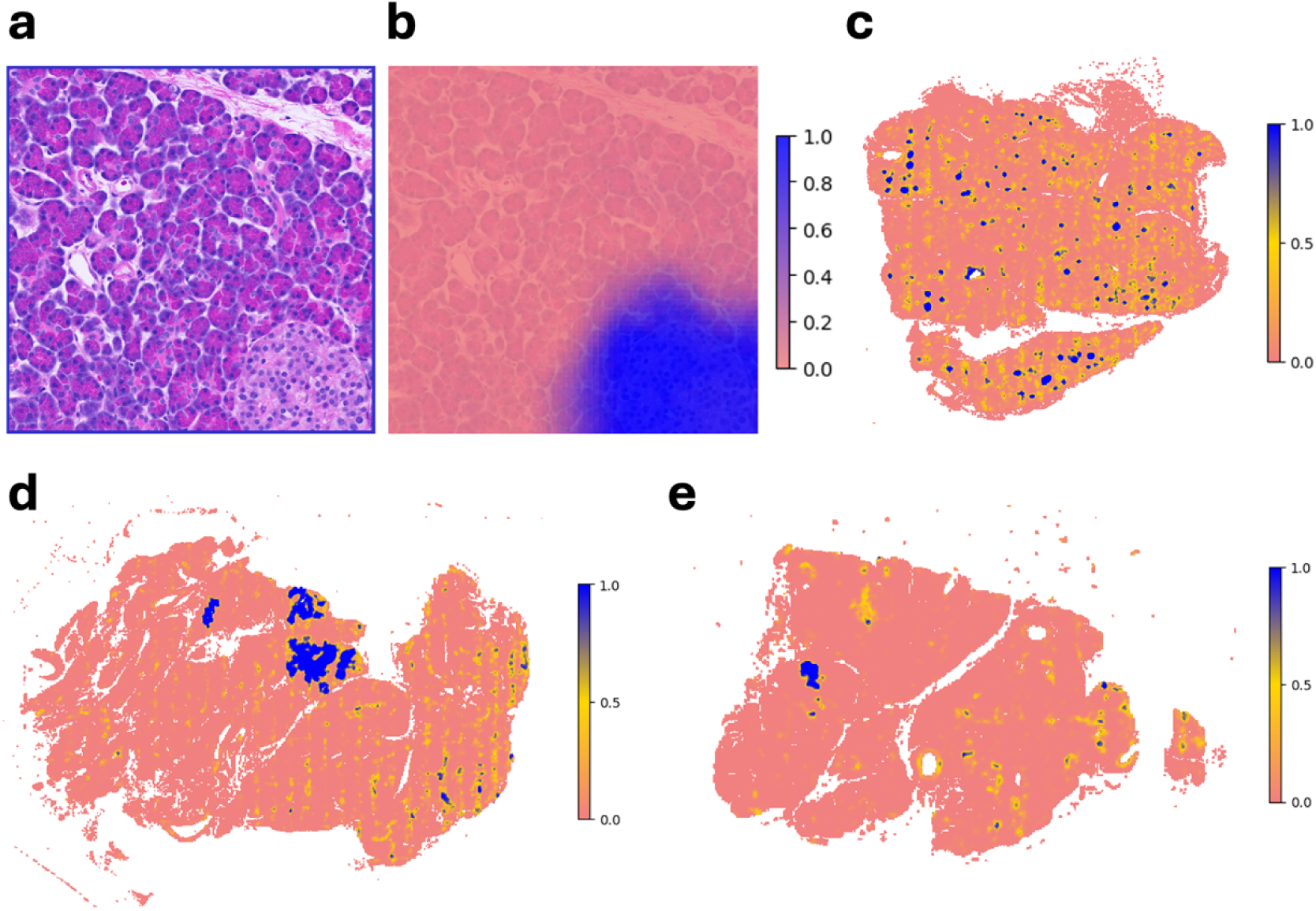
Spatial inference of islets and neoplasms in human pancreas. **a.** Example image patch from the pancreas of a 23-year-old female donor, with an islet in the lower right. **b**. The inferred islet spatial probability distribution within this patch. The classifier predicts a 0.94 probability that the patch contains an islet**. c.** Distribution of predicted islets in the complete pancreatic tissue sample. Coloring indicates predicted islet probability. **d.** pancreas head superior and **e.** head inferior from a 69-year-old female donor. Coloring indicates predicted probability of PanIN.

We next trained a classifier for pancreatic intraepithelial neoplasia (PanIN), a recognized precursor to pancreatic ductal adenocarcinoma. We interactively annotated 15 positive and 25 negative patches (total tissue size: 2063 patches) from the head superior and head inferior regions of the pancreas from a 69 y.o. female donor (**Fig. 3d,e**). In both samples, extensive PanIN regions were identified, with strong validation on pathologist annotations (AUROC 0.996, AUPRC 0.633). Pathologist-verified pixel fractions of PanINs in these slides were 1.6% and 0.5%, respectively, both considerably lower than the AUPRC. To test the robustness of the classifier, we applied it to 3 tissue sections from the 23 y.o. and 71 y.o. donors, again achieving excellent validation with average AUROC 0.99, AUPRC 0.376, and FPR 0.0026. We further analyzed 5 tissue sections from these donors prepared by a different protocol, i.e. Xenium processing prior to H&E imaging. We trained a classifier for these samples by interactively annotating 9 positive and 114 negative patches (total tissue size: 1524 patches), yielding a classifier with average AUROC 0.99, AUPRC 0.567, and FPR 0.0005.

### Interactive classification of kidney and fetal membranes spatial structures

We interactively trained classifiers for multiple kidney histological structures, including glomeruli, obsolescent glomeruli, and fibrous tissue, as in **Supplementary Fig. 5.** First, we trained a classifier to detect glomeruli in 4 kidney tissue sections spanning regions of renal cortex, medulla, and pelvis by interactively annotating 42 positive and 42 negative of 2884 total patches. We achieved average AUROC/AUPRC of 0.98/0.77. To detect obsolescent glomeruli we annotated 16 positive and 58 negative patches and achieved AUROC/AUPRC 0.93/0.16 and FPR 0.017. This AUPRC value, while low on an absolute scale, is much higher than the prevalence of obsolescent glomeruli in this tissue, which cover only 0.002 of the pathologist annotations on average. To detect fibrous tissue, we annotated 30 positive and 30 negative patches from 11356 total patches, achieving average AUROC/AUPRC values of 0.9/0.85. Here we used 4×4 tile patches because the wide prevalence of fibrous tissue required a higher resolution approach. Generation of these classifiers was completed in only a few minutes of interactive training. A demonstration of the histology interactive workflow for kidney fibrous tissue is provided in Video S1.

We next applied DIANNE to layered fetal membrane samples from a 34-year-old patient, **Fig. 4**, Panels (a)–(c). We interactively trained classifiers for red blood cell (RBC) aggregates, amniotic membrane, and chorionic membrane, each of which is important for fetal membranes pathology, where membrane integrity and membrane stromal features can indicate gestational abnormalities or maternal-fetal complications [29,30]. DIANNE was effective for each of these, as validated by comparison to pathologist annotations. Positive training patches / negative training patches / AUROC / AUPRC were 16/16/0.97/0.72 for RBC aggregates, 30/30/0.95/0.72 for amnion, and 21/21/0.95/0.86 for chorion.

**Figure 4.**
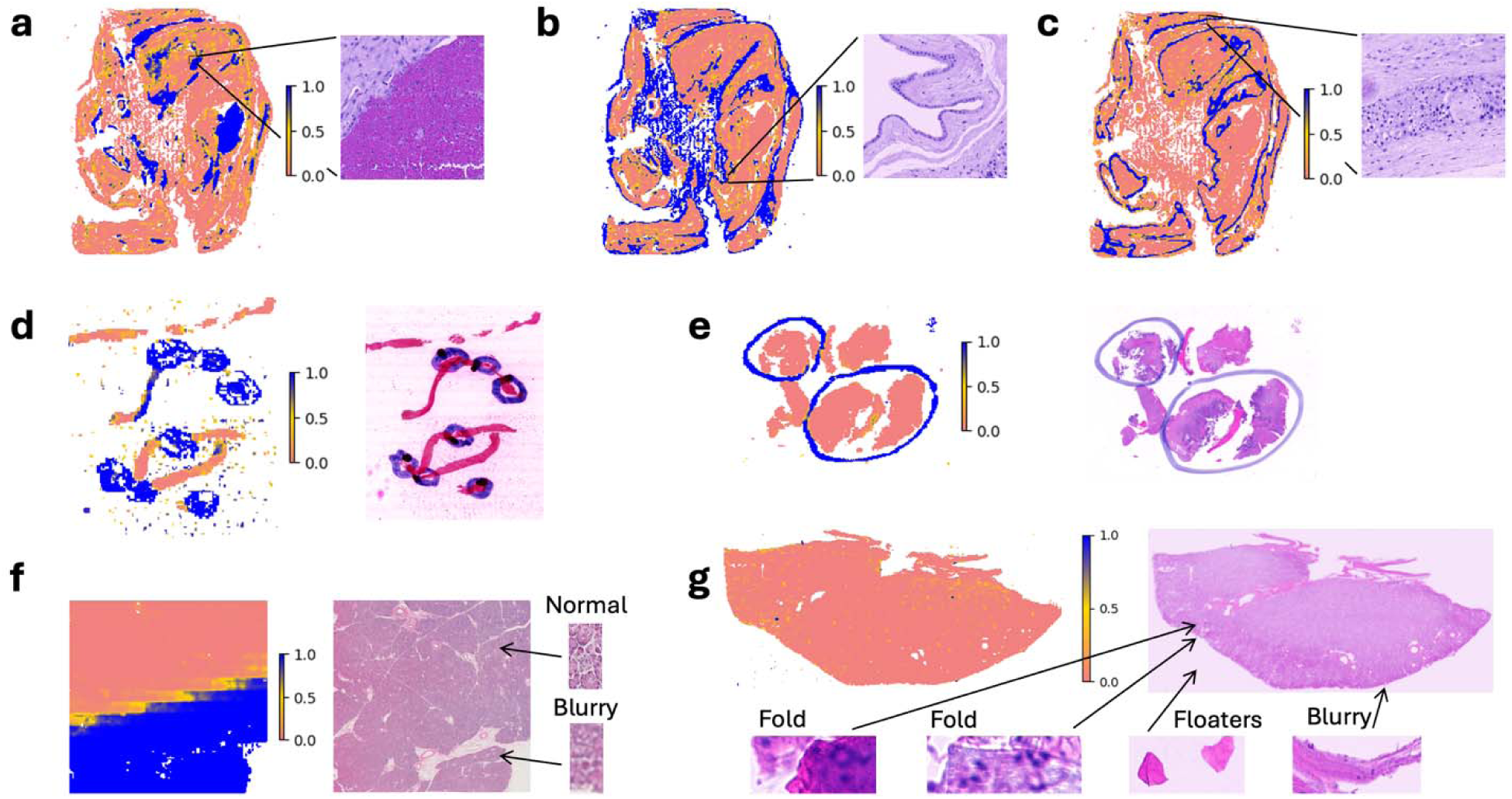
Spatial classifiers for fetal membranes and tissue artifacts. In all panels, the right side shows a zoomed-in example H&E region while the left side shows classifier predictions over the whole tissue section. **a,** Spatial inference of red blood cell aggregates, **b,** amniotic membrane, and **c,** chorionic membrane, all from human membrane tissue from a 34 y.o. patient. **d**, Pediatric sarcoma H&E slide with blue ink markings. The ink markings and background stain droplets are classified correctly as artifacts by the interactively trained classifier. **e**, Another pediatric sarcoma H&E slide with blue ink markings. **f,** WSI of a normal pancreatic tissue with extensive out-of-focus regions, identified as artifacts (blue). **g,** WSI of a human kidney with only minor staining artifacts, noted in the insets.

### Interactive classification of image artifacts

Classifiers for image artifacts such ink markings and background stain droplets (**Fig. 4d and 4e**, pediatric sarcoma slides) could also be accurately trained from small numbers of labeled patches. Classification metrics (positive training patches / negative training patches / AUROC / AUPRC) were 5/5/0.95/0.85 and 10/10/0.99/0.94 for ink markings and background stain droplets, respectively. Classifiers for out of focus regions were also accurate (**Fig. 4f**, pancreatic tissue slide), with positive/negative/AUROC/AUPRC values of 5/5/0.99/0.99. Slides with only minor pathologist-identified artifacts were successfully classified as normal, e.g. in a kidney slide shown in Fig. 4g. We trained a classifier for artifactual regions from this kidney slide, yielding positive / negative / AUROC /AUPRC = 6/109/0.96/0.48. Thus DIANNE can effectively support interactively-trained quality control for artifacts.

### Application of DIANNE to IHC images

DIANNE also provides capabilities for training classifiers using immunohistochemistry (IHC) foundation models. For example, we analyzed tiles from patient 059 from the ACROBAT dataset [33] of HER2-stained breast cancer, which we clustered into eight distinct classes using UNI2 [15] image features. A clinical pathologist annotated this slide at pixel resolution and confirmed the clustered classes correspond to meaningful categories including invasive carcinoma with varying HER2 expression, DCIS, fibroadipose tissue, and adipose tissue. We interactively trained classifiers using this IHC slide with PCMA augmentation parameter 0.5 (**Supplementary Fig. 6**) and validated using the pathologist annotations. We considered four types of regions and observed strong performance for each: non-tissue background (AUROC/AUPRC/recall 0.99/0.96/0.89), HER2-negative adipose tissue (AUROC/AUPRC/recall 0.97/0.95/0.99), DCIS with HER2 2+ staining (AUROC/AUPRC/recall 0.99/0.68/0.84), and invasive carcinoma tissue (AUROC/AUPRC/recall 0.96/0.82/0.80). For comparison, we trained non-PCMA classifiers for the same tasks and the same annotated patches. The non-PCMA classifiers also performed well, but tended to have lower recall: background regions (AUROC/AUPRC/recall 0.97/0.94/0.70), HER2-negative adipose tissue (AUROC/AUPRC/recall 0.98/0.99/0.96), DCIS with HER2 2+ staining (AUROC/AUPRC/recall 0.98/0.50/0.81), and invasive carcinoma (AUROC/AUPRC/recall 0.96/0.85/0.76). All four of the PCMA-trained classifiers converged rapidly in the interactive training (**Supplementary Fig. 6e-h**). None used more than 15 positive or 15 negative annotated patches. Remarkably, the tissue background classifier achieved its excellent performance (AUROC/AUPRC/recall 0.99/0.96/0.89) with only a single positive and a single negative training patch.

### Application of DIANNE to multiplex spatial proteomics images

DIANNE also enables classifier training based on multiplex spatial proteomics data. This is done by integration of the KRONOS multiplex protein foundation model [17] with a visualization tool that converts mIF data to RGB images for interactive patch labeling. The multiplex spatial proteomic visual annotation workflow is shown in **Fig. 5**. Panel (a) shows the corresponding Jump-start tool, which allows users to convert multiplex proteomics data into RGB images, view representative patches, and initiate the first positive patch. For example, in **Fig. 5** we took as input 39 channels of PhenoCycler Fusion data from two tissue sections. We converted these channel data into RGB for visualization, though we note that the classifier uses all channels for training. We chose the rightmost patch in **Fig. 5a** for initiation of training for an PanIN classifier, as this patch contains papillary tissue that is characteristic of PanIN. **Fig. 5b** illustrates a new proposed patch for iterative curation. The updated spatial probability map for PanIN is shown (**Fig. 5c**) to aid visual evaluation. We annotated 11 positive and 39 negative of 374 total patches (patch size 6×6 tiles, tile size 112 mm, image resolution 0.5065 mm per pixel). The resulting classifier achieved AUROC 0.995, AUPRC 0.867, FPR 0.024 at probability threshold 0.9 for the first tissue section, with the true prevalence of PanIN being 1.8% (**Fig. 5d**). In the second tissue section, we found AUROC 0.923, AUPRC 0.283, FPR 0.0004 at probability threshold 0.9, with true prevalence 0.28% (**Fig. 5e**).

**Figure 5.**
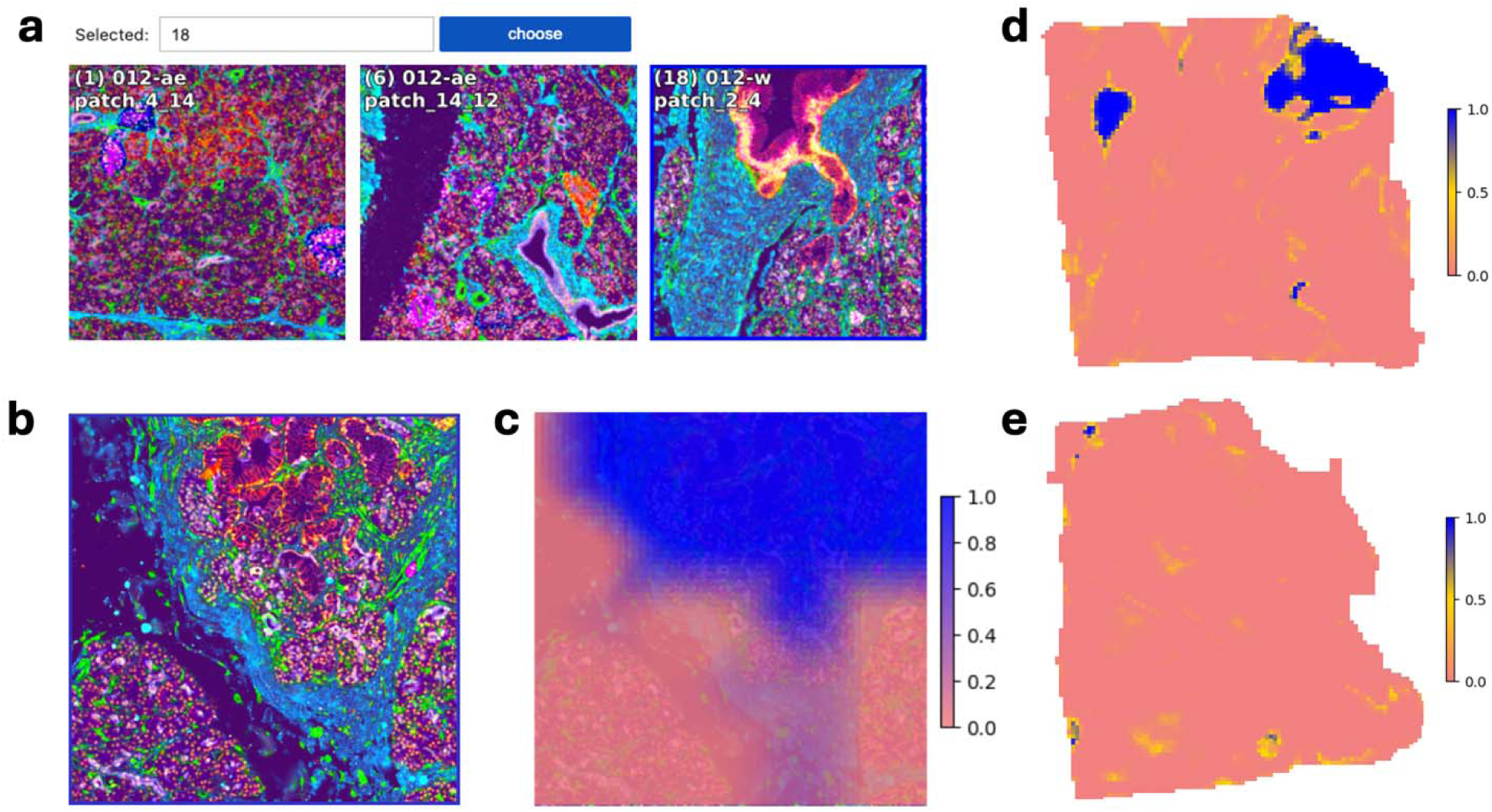
DIANNE analysis of PhenoCycler Fusion 39-channel spatial proteomics data from a 69 y.o. human pancreas donor. **a**, The DIANNE Jump-start tool converts the multiplex proteomics data to a 3-channel RGB image and displays representative image patches. The operator has selected the rightmost patch, outlined in blue, to initiate training. **b**, Example of a subsequent patch displayed for operator curation. **c**, Probability map for the current PanIN classifier in the same patch as **b**. **d** and **e** show probability inferences from the final classifier for PanIN in **d,** pancreas head superior and **e,** pancreas head inferior sections, respectively.

### Analysis of H&E+Spatial Transcriptomic Data within the Molecular Static Workflow

The molecular static workflow enables training based on a fixed set of H&Es with matching spatial molecular data. The workflow uses transcriptomic behaviors in tiles to distinguish patches for training the H&E-based classifier. The annotations may be chosen by the user to be based on cell type counts or gene expression. For example, we trained a classifier for islets (**Fig. 6**) using Xenium *in Situ* profiling + H&Es in 30 human pancreatic tissues, then tested on tissues from two donors—23-year-old donor 008 and 71-year-old donor 002. Panels (a) and (b) illustrate tile-level counts of endocrine cells. Training is done at the patch level (augmentation parameter a=0.95), and DIANNE assesses patches (8×8 tiles) as positive when more than 75 endocrine cells are present. For validation, we used the Xenium data to define true islet tiles as those with >=10 endocrine cells. Metrics of islets inference on donor 008 (panel a, JDC-WP-008-v) were AUROC 0.955, AUPRC 0.520, FPR 0.003 at probability threshold 0.9, true prevalence 2%. For donor 002 (panel b, JDC-WP-002-v) these were AUROC 0.935, AUPRC 0.476, FPR 0.003 at probability threshold 0.9, true prevalence 3%.

**Figure 6.**
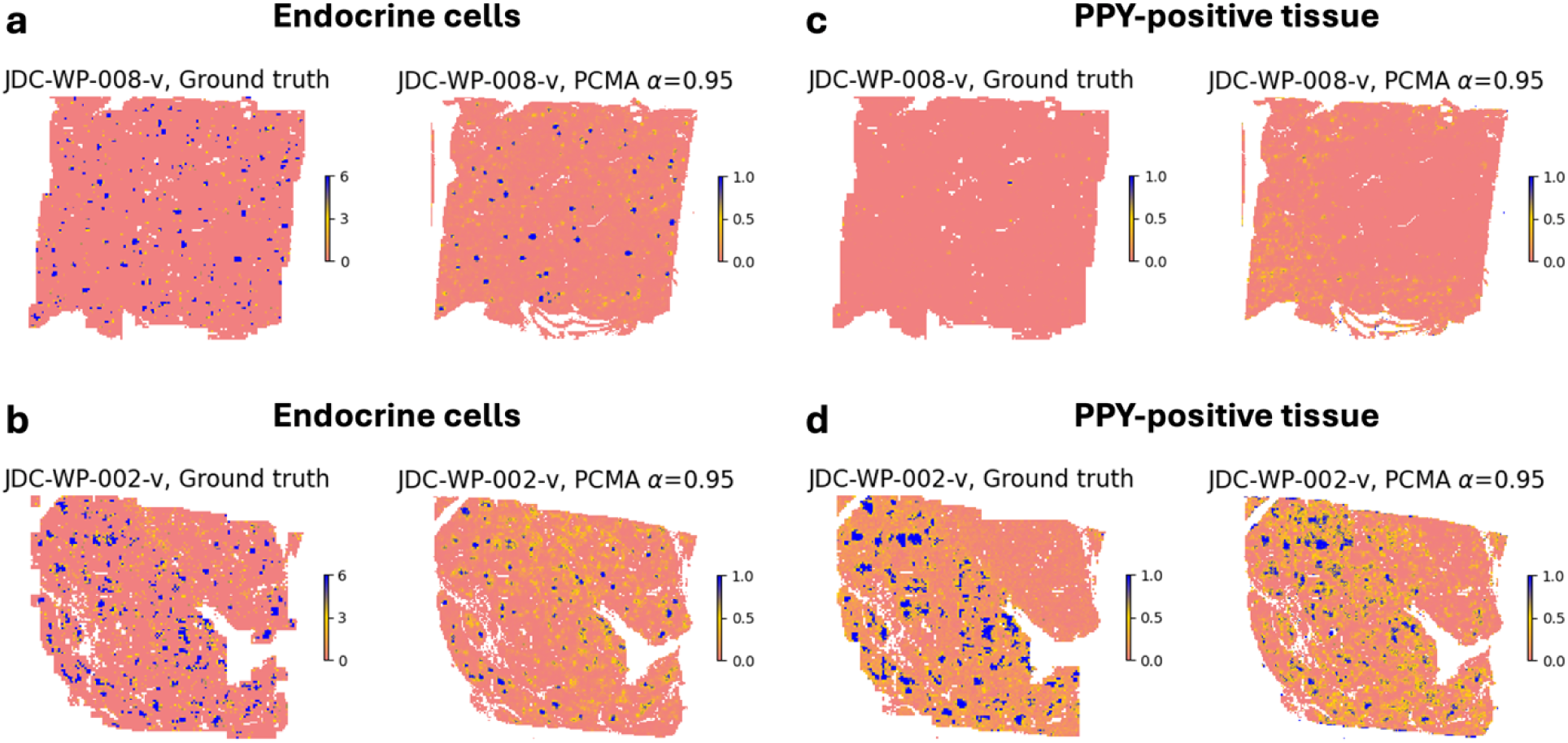
Xenium-based data and classifier inferences for human pancreases (23 y.o. donor 008 and 71 y.o. donor 002). **a** and **b:** (left) Xenium-based tile-level counts of endocrine cells. Tiles with 6+ cells are shown as 6. (right) DIANNE spatial inferences from an H&E-based classifier trained via a patch threshold of 75 endocrine cells. **c** and **d:** (left) Xenium-based tile-level average PPY expression. (right) DIANNE spatial inferences from an H&E-based classifier trained via a Xenium patch threshold of PPY > 0.2.

We also trained a classifier for PPY expression, a gene associated with gamma cells in islets in healthy individuals, using the same train and test sets as above [34]. Panels (c) and (d) show tile-level average PPY expression, with regions assessed as positive when average PPY expression > 0.2. We did not observe PPY-positive tissue in donor 008 (panel c) but for donor 002 (panel d) the metrics were AUROC 0.821, AUPRC 0.407, FPR 0.006 at probability threshold 0.9, corr. 0.346, prevalence 11%.

### Training of H&E+Spatial Transcriptomic Classifiers with the Molecular Interactive Workflow

The molecular interactive workflow enhances the histology interactive workflow by integrating transcriptomics data into the patch view. Importantly, during iterative training the workflow retrieves the Xenium Zarr-stored transcriptomics data only for the current patch. This selective retrieval allows transcripts, cell boundaries, and annotations to be loaded and rendered within seconds for display atop the histological image (either H&E or antibody-stained). The result is a responsive and scalable interface for real-time exploration of spatial gene expression patterns with tissue morphology.

We illustrate the molecular interactive workflow for PanIN classification in **Fig. 7**. A corresponding video demonstration is provided in **Video S3**. Here we hypothesized MUC5AC patterns might clarify PanIN beyond H&E-only labeling, as MUC5AC is known to be associated with PanIN but their spatial correlations have not been well-quantified. This example shows local MUC5AC expression in human pancreatic tissue displayed for user assessment during iterative training. Panel (a) shows the DIANNE toolkit interface, where Xenium-derived MUC5AC transcripts are visualized as green dots overlaid on a patch. Panel (b) presents a tile-level probability heatmap, interpolated for smoothness. Regions with higher predicted likelihood of the MUC5AC pattern are colored blue, aiding the user in making informed decisions during patch-level classification. To train the classifier, we interactively annotated 16 positive and 24 negative of 770 total patches, achieving AUROC 0.991, AUPRC 0.897, Pearson correlation coefficient 0.677, and FPR 0.005 at probability threshold 0.9. The ground truth for AUROC/AUPRC/FPR was binarized at a threshold of 20 transcripts per tile. Panel (c) shows the whole-slide, revealing high-probability PanIN regions predicted by the classifier. This transcript-informed classifier is more accurate than the H&E-only trained classifier for the same slides (**Fig. 3**, AUPRC 0.567). It is also superior to the mIF PhenoCycler-based classifier (avg AUROC 0.959, AUPRC 0.575, FPR 0.012).

**Figure 7.**
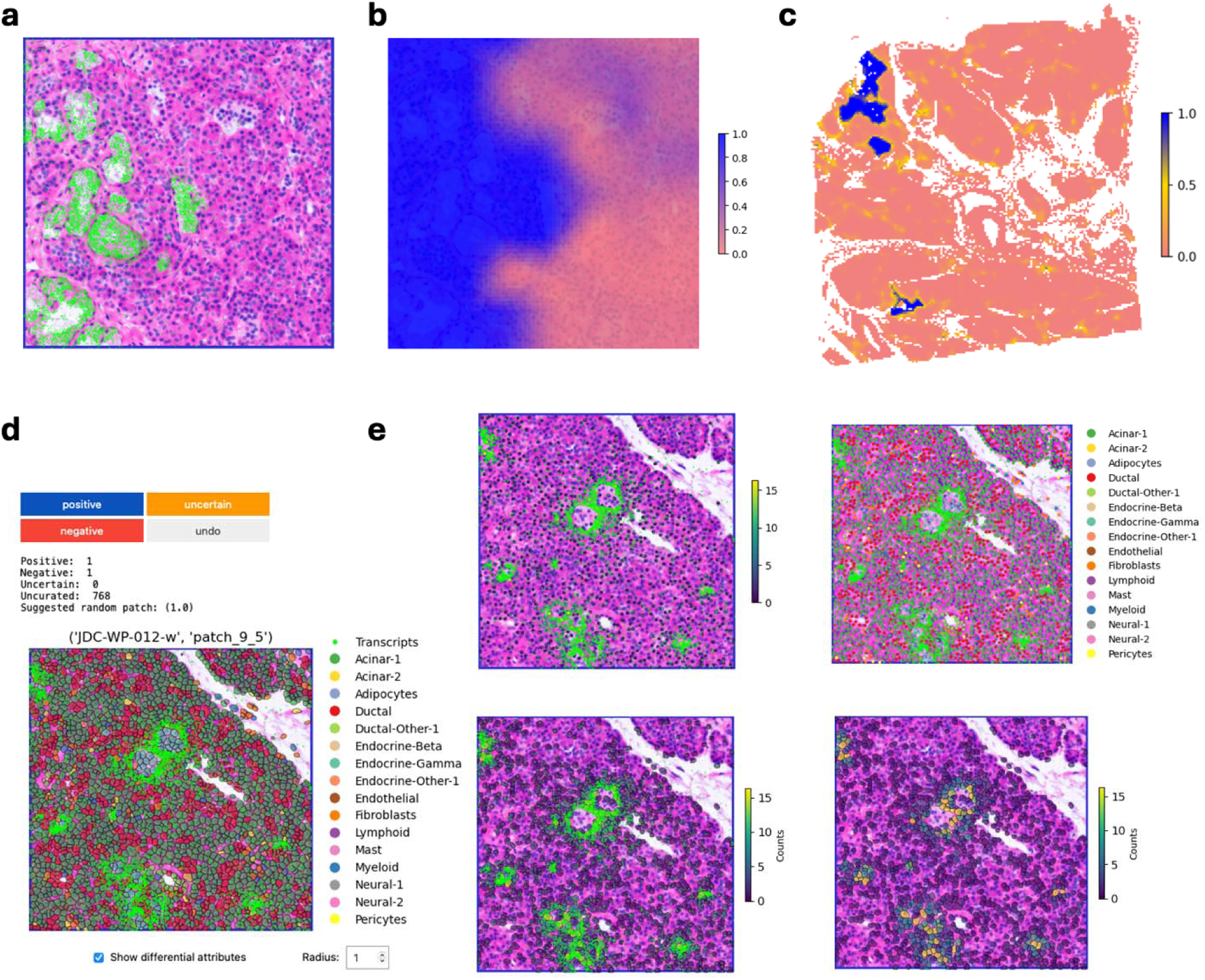
Illustration of the molecular interactive workflow. Xenium transcripts of MUC5AC are used to enhance interactive annotation of PanIN patches from an H&E of a human pancreas from a 69 y.o. donor. **a**, DIANNE Jupyter view showing Xenium transcripts as green dots overlaid on the H&E patch. **b**, The current tile-level positive class (MUC5AC+ PanIN) probability heatmap during interactive classification, overlaid on the same image patch. **c**, Whole slide probability heatmap denoting high MUC5AC+ PanIN probability regions (blue) after completion of interactive training. **d.** Visualization of molecular data on H&E images, including cell boundaries, cell types, and Xenium transcripts of PPY, GCG, INS, SST. All transcripts are displayed in green. **e**, Alternative display of cells as either circles (top two panels) or segmented objects (bottom two panels). Cells are color-coded according to gene expression (left) or cell annotation labels (right).

DIANNE provides versatile molecular data visualization within the annotation interface, enabling consideration of both gene expression and cell-type. In Panel (d), Xenium transcripts for PPY, GCG, INS, and SST on H&E-stained images are shown alongside cell boundaries and user-specified cell type annotation labels, providing a multi-modal view of the tissue microenvironment. Panel (e) highlights the flexibility of the visualization settings, where cells can be rendered either as simple circles or as segmented objects, with color-coding based on gene expression levels or other annotations. This configurability enhances the user’s ability to take into account complex spatial relationships and molecular signatures during classifier training. DIANNE is also applicable to 10x Atera data, as shown for a public in situ breast cancer dataset in **Supplementary Figure 7**. The freehand GUI integrates the H&E image, mIF staining, and spatially aligned cell annotations and transcripts. Classifiers were trained from freehand annotations of Atera data for tumor cells, basal-like DCIS cells, invasive carcinoma cells, and lymphocyte-infiltrated stromal regions, illustrating DIANNE’s capacity to enable rapid, annotation-driven spatial classification.

## Discussion

DIANNE introduces Positive Class Mixup Augmentation (PCMA) to enable rapid, flexible and accurate spatial classifier training. We have demonstrated the effectiveness of DIANNE for key types of spatial classification, including cancer detection, identification of diverse tissue-specific structures, and artifact removal for H&E, IHC, mIF, and spatial transcriptomic inputs. Rather than training directly on positive and negative examples, DIANNE mixes positive and negative samples into augmented positive samples. This provides an effective regularization that enables accurate classification development from only a small amount of training.

Importantly, the speed of DIANNE facilitates exploratory analysis, empowering users to interactively train classifiers via expert knowledge, concordant with pathology practice. Prior methods have typically hard-coded parameters that should be structure-specific, such as the threshold fraction for a tile to be considered positive [35]. In contrast, DIANNE emphasizes flexible testing, evaluation, and revision, which are central to the discovery process. The speed needed for these capabilities arises from the efficiency of the SAMPLER representation. Notably, SAMPLER yields performance superior to moment-based representations, even though they also compactly encode distributional information. This is likely because the natural ordering of quantile features makes them well-suited to logistic regression-based classification.

DIANNE advances prior human-in-the-loop approaches [36] by leveraging large pathology foundation models for tile-level feature extraction to capture distributional tissue context. For comparison, nuclei.io recently developed an active learning with human feedback system to speed nucleus classifier development based on prioritization of nuclei for pathologist review [37], and [38] used foundation model fine-tuning for kidney nuclei segmentation. DIANNE handles the more general patch-level problem, not restricted to nuclear morphology. As another comparison, HistoROI developed a human-in-the-loop patch classifier by prioritizing internally homogeneous patches within WSIs for pathologist annotation into known morphological categories [39]. DIANNE, on the other hand, handles intrapatch heterogeneity quantitatively via the SAMPLER representation, providing fundamental advantages in what data can be considered. More broadly, DIANNE’s approach for exploration of novel structures can be considered as a type of online learning, a powerful interactive human-in-the-loop paradigm that has been limited by real-time compute capabilities [40]. DIANNE addresses this prior limitation via the SAMPLER representation and PCMA, which are critical to its effectiveness.

DIANNE provides capabilities for IHC and multiplex protein images using corresponding foundation models, which are important for clinical considerations such as HER2+ breast cancer [41]. These workflows support the discovery of protein spatial behaviors, e.g. immune cell markers or gradients of signaling molecules. Because H&E, IHC and spatial proteomics foundation models are rapidly developing [42,43,44], the DIANNE framework allows modular swapping of foundation models. DIANNE also has capabilities for both static and interactive analysis of spatial transcriptomics data. Our current approaches use H&Es co-registered to spatial transcriptomics data along with H&E foundation models, since spatial transcriptomics foundation models are not yet standardized. These are being actively developed [45,46,47], but the high dimensionality and complexity of spatial transcriptomics data have hindered the definition and acceptance of ground truth sets. We believe that DIANNE can play an important role in addressing this challenge, by providing a means for pathologists and researchers to expertly define, evaluate, and annotate gold standard sets.

Overall, DIANNE can assist scientists in working with diverse multimodal foundation models to discover, define, annotate, and predict new spatially important structures in tissues. Integrating diverse biological data types—such as imaging, transcriptomics, and proteomics—into unified models enables deeper insights into cellular mechanisms and tissue architecture [48]. DIANNE provides an efficient and accurate framework for exploring and training classifiers for accelerating discovery in spatial biology.

## Methods

### Spatial Inference Workflows

We implemented four distinct workflows—histology static, histology interactive, molecular static and molecular interactive—that incorporate PCMA during slide- or patch-level classifier training using SAMPLER-based image representations. These workflows operate with the Spatial Transcriptomics Quantification (STQ) pipeline [12] for H&E or the Multiplex Immunofluorescence Extraction (MIE) pipeline from Spatial Omics Tools (SOT) output for multiplex immunofluorescence images.

The STQ pipeline processes WSIs in an arbitrary grid configuration with either user-provided or de novo generated tile coordinates. It performs slide normalization, tile grid creation, and imaging feature extraction using histopathology foundation models. STQ implements current deep convolutional neural networks (CNN) or vision transformer (ViTs) foundation models that have been pre-trained on large-scale histopathology datasets. Examples include CTransPath [13] for all H&E slides and UNI2 [15] for IHC data. The normalization step ensures consistency across slides and mitigates batch effects. Similarly, the MIE pipeline processes mIF WSIs to extract tile-level imaging features using a spatial proteomics foundation model, KRONOS [17]. Tile features provide inputs for classifier training and spatial inference.

To accommodate diverse analytical needs, DIANNE provides a modular workflow architecture flexible for four binary choices: static versus interactive execution, slide-level versus patch-level analysis, histology versus histology+molecular data, and spatial versus non-spatial molecular data (**Supplementary Fig. 8a**). This set of choices is illustrated in **Supplementary Fig. 8a** in orthogonal planes. Histological workflows can be based on either hematoxylin and eosin (H&E) or multiplex immunofluorescence antibody-based staining, while molecular workflows support both spatially resolved and bulk data types (**Supplementary Fig. 8b**). Workflow configurations are indicated with checkmarks or circles, providing a simple interface for analyses.

Histology static utilizes either slide- or patch-level SAMPLER representations of imaging features for classifier training. Each WSI is labeled as either positive or negative for a target attribute (e.g., cancer presence, tissue artifacts). In patch-level representations, the patch size is chosen such that patches both with and without the desired histology attribute exist; by default, we use 8×8 tiles. This approach enables straightforward training and spatial classification in the absence of spatially refined annotations. PCMA is applied during training to enhance feature representation. A summary is illustrated in **Fig. 1**, and slide- or patch-level flowcharts are shown in **Fig. 8a and 8b**, respectively. Definitions of slide, patch and tile are shown in **Fig. 8c**.

**Figure 8.**
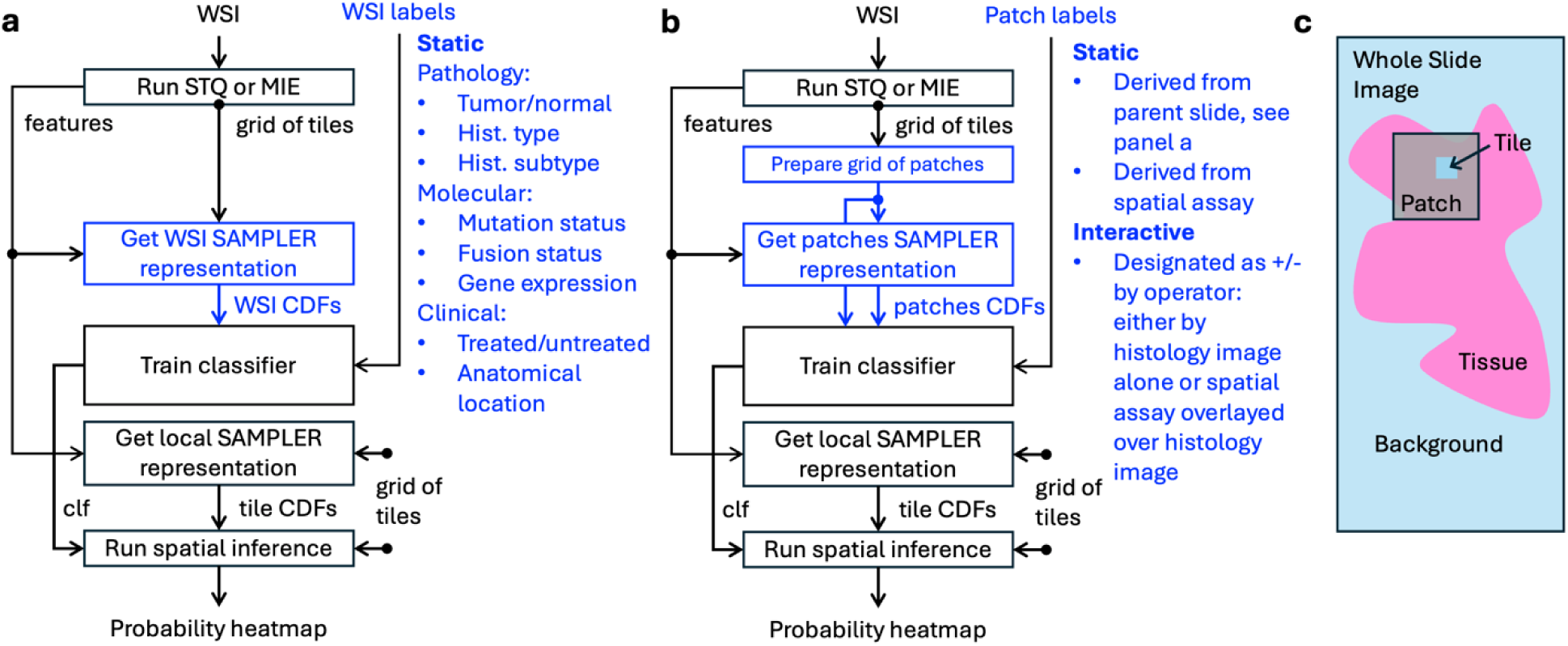
Two categories of workflows for spatial SAMPLER inference. Workflows process WSIs with STQ (H&E and IHC images) or MIE (mIF images), make a grid of tiles, and extract imaging features for each tile. This yields a spatially resolved differential attribute probability heatmap. All workflows use PCMA during the ‘Train classifier’ step. **a**, In WSI-based workflows, slides’ SAMPLER representations are directly used for classifier training, based on weakly supervised (slide-level) training for a target attribute. **b**, Patch-based workflows operate on patches of tiles (default 8×8). **c**, Schematic of tiles, patches, and slides. All interactive workflows are patch-based and display test patches during the interactive annotation process. Differences between the WSI- and patch-based workflows are highlighted in blue.

The histology interactive workflow introduces spatially refined annotations by operating on patches of tiles (typically 8×8, adjustable). During interactive annotation, patches are displayed to the operator using an annotation toolkit. Patch selection is guided by an active learning algorithm informed by the current classifier state, optimizing annotation efficiency and classifier performance.

The molecular static workflow integrates spatially resolved molecular data with histology images. It operates on tile patches and uses molecular data (e.g., from 10x Visium, Xenium In Situ or Atera In Situ assays) to derive patch-level labels without displaying raw imaging or molecular data during training. When using 10x Visium data, the output grid can be directly supplied to the STQ pipeline.

The molecular interactive workflow combines histology interactive and molecular static workflows, enabling overlay of molecular data on histology patches during annotation. This integration supports more accurate and informed labeling, particularly when high-resolution spatial transcriptomics data are available.

### Graphical user interface

We developed two complementary labeling interfaces, Guided and Freehand, to accommodate different annotation workflows. In the Freehand interface, **Fig. 1a**, the user draws positive or negative contours directly onto the image, and all patches falling within the drawn region are labeled automatically. Unlike the Guided interface, freehand annotation is performed on one slide at a time, allowing the user to freely navigate the image — zooming, panning, and selecting regions of interest based on their own visual judgment rather than algorithm-driven patch proposals. This gives the user full spatial control, making it particularly valuable when the user has strong domain expertise and wishes to directly target specific tissue structures or regions. However, annotations accumulated across all slides are pooled together for model training. In the Guided interface, **Fig. 1b**, patches are presented to the user one by one, and the user designates each as positive or negative using dedicated buttons. Patches are drawn simultaneously from across all loaded whole slide images via a patch proposal algorithm, **Supplementary Fig. 1**, ensuring balanced and representative sampling across the entire dataset. This approach is well-suited for careful, systematic review of ambiguous or fine-grained regions, ensuring no patch is overlooked. Together, the two interfaces allow users to balance annotation speed and precision according to the complexity of the image content, and both support multi-slide workflows that aggregate labeled patches into a unified training set.

### Data augmentation with PCMA

The SAMPLER representation for an image is computed by first calculating foundation model deep learning features in all tiles and then, for each feature, calculating the distribution of feature values across all tiles [11]. Classifiers can be trained using logistic regression on positive and negative samples with the SAMPLER representation. DIANNE improves on this procedure by using positive class mixup augmentation (PCMA) during classifier training. PCMA replaces the SAMPLER representation for each positive sample with an improved representation informed by negative samples. The procedure is: 1) For each positive sample (WSI or patch), randomly pick one negative sample. 2) Calculate the SAMPLER representations for the positive sample and for the negative sample. 3) Integrate the positive sample and negative sample representations in a weighted fashion (parameter a) to yield an augmented positive sample. 4) Replace the SAMPLER representation for the positive sample with the augmented SAMPLER representation, 5) Repeat steps 1-4 for each positive sample. 6) Retrain the classifier.

Importantly, each positive sample is augmented by a single contiguous negative sample, rather than by an ensemble of non-contiguous negative tiles. This contiguous sample-based approach better maintains spatial distributional relationships and improves classification accuracy. We note that sample augmentation is only used during classifier training. Trained classifiers are applied to unaugmented data during spatial inference.

The weighted integration of a positive and negative sample (i.e. step 3 above) is done as follows. The SAMPLER representation is based on the distributions of the deep learning feature values across tiles, approximated using the deciles of the cumulative distribution function (CDF) for each deep learning feature [11]. We developed a function (getDiscreteCombinedCDF) to integrate the positive sample CDFs and negative sample CDFs without having to re-access the original tile feature values, enabling rapid data augmentation. It takes as input parameters the quantile and feature values for each of the two input CDFs, as well as weight parameter α. The function calls a supplementary function getDiscretePDF that extrapolates the CDF to a range [0, 1], generates the continuous probability distribution functions (PDF), interpolates it to 200 points, and integrates them to get discrete PDFs, *p_1(x_1_ ∈ S_1_)_* and *p_2(x_2_ ∈ S_2_)_* where *S*_1_ and *S*_2_ are the sets of tile feature values from a positive and a negative sample, respectively. Then it generates a discrete PDF interpolated into N points from each of the two coarse CDFs. The resulting discrete PDF values for each feature are concatenated and sorted by the feature value. The cumulative weighted sum of the values is then computed from sorted concatenated discrete PDFs:

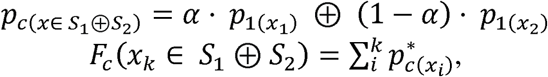

where * indicates the sorting operation ⊕, indicates a mathematical concatenation operation. Since each of the discrete PDFs sums to one, *p_c(x_1_ ∈ S_1_ ⊕ S_2_)_* is a valid normalized probability distribution function, from which the fine-grained CDF,*F_c_*, is computed. The CDF values are interpolated to match the range of the input quantiles *{0.05(1 + k) | k ∈ {1,2,…,9}}* to obtain *F∧^*^_c_(x)*

For example, in **Supplementary Fig. 9a**, the combined CDF,*_a_* is generated by concatenating the feature values from the two samples and computing the quantiles. The combined *CDF_b_* is generated by naively combining the two CDFs in a tile-based fashion with weights α = 0.74, based on the relative tile counts from the tumor and normal samples α(1 − α) = 2.8. The combined CDF,*F_c_* is generated by combining the two CDFs with weights α = 0.5. It is useful to compare the behaviors of different deep learning features. As can be seen in their CDFs, the feature 135 values of the tumor sample, *CDF_2_*, are generally larger compared to the normal sample, *CDF_1_*. Feature 135 has a larger difference in the CDFs between the two samples compared to feature 15, see **Supplementary Fig. 9b**. In the combined CDF, the smaller quantiles of feature 135 are dominated by the normal sample, while the larger quantiles are dominated by the tumor sample. **Supplementary Fig. 10** shows CDFs at different values of the mixing parameter. For close to 0 (violet), there is little mixing of the negative class into the positive class, while large values of close to 1 correspond to red. Edge cases approach either *CDF_1_* or *CDF_2_*.

Because generation of *CDF_a_* requires loading and processing higher volume of data than for *CDF_b_*, the effective speedup of generating *CDF_b_* over *CDF_b_* is up to 100-fold, depending on the sizes of WSIs in the dataset.

### Patch selection algorithm

The algorithm, **Supplementary Fig. 1**, starts by randomly generating a number in the interval [0,1). If this number exceeds a predefined threshold, e.g. 0.35, the algorithm checks whether at least one positive-labeled and one negative-labeled patch have been curated. When such labels exist, a classifier is trained on the curated data and applied to infer positive class probabilities for all remaining uncurated patches. The algorithm then proposes patches strategically: if more positive patches are already curated, it selects the most-likely negative uncurated patch (the one with lowest predicted probability); otherwise, it selects the most-likely positive uncurated patch (the one with largest predicted probability). We select confident rather than uncertain patches, aiming to capture features that define the positive class, rather than refine the classifier’s decision boundary. If the random number falls below the threshold, or if positive and negative labels do not yet both exist, the algorithm simply proposes a random uncurated patch, ensuring that the algorithm does not stall in local minima. Each proposed patch is then curated by the user, after which the algorithm checks whether uncurated patches remain and repeats the process accordingly. This iterative procedure continues until all patches have been curated, at which point the algorithm terminates. The threshold parameter controls the balance between exploration through random sampling and exploitation through model-guided active learning, with higher thresholds promoting greater variety in the proposed patches.

### Normal tissue data selection

A total of 30,105 SVS file names were retrieved using Google BigQuery from the TCGA dataset (slide_images_gdc_current). Files corresponding to tissue blocks labeled 11A were retained, resulting in 2,701 entries. To ensure consistency in image quality, additional filtering was applied: only images with a resolution between 0.24 and 0.26 μm/pixel were included, yielding 503 files. Finally, files with sizes between 30 MB and 230 MB were selected, resulting in 305 files representing 310 tissue images. A total of 20 normal lymph node slides were downloaded from CAMELYON16 and split into 31 tissue images.

### Export GeoJSON contours from DIANNE probability maps

The inferred probability heatmaps are exported to GeoJSON format compatible with QuPath using a dedicated DIANNE tool ‘extractContoursForQuPath’. First, a whole slide probability map is optionally smoothed using a Gaussian blur (sigma 6.25) and binarized with an adjustable threshold (default 0.5). Contours are extracted using OpenCV with RETR_TREE and CHAIN_APPROX_SIMPLE, and those below 100 pixels in area are discarded. Contour coordinates are rescaled to the original image space by the downsampling factor. Holes are preserved as inner rings of GeoJSON Polygon features, with nested sub-regions recursed as independent top-level features. The resulting GeoJSON feature collection is saved as a lightweight text file, with each feature carrying QuPath-compatible annotation properties including class label and display color, allowing the exported annotations to be directly imported into QuPath for visual validation of model predictions and downstream spatial analysis.

### Pathologist slide annotations for classifier validations

A pathologist manually annotated selected slides in QuPath 0.5.1 at the pixel level. Each of these manual annotations was subsequently converted into whole slide pixel masks and aggregated at the tile level to evaluate performance and validation metrics. The slide annotations are detailed in **Supplementary Table 1** and encompass the following:

- Sarcoma: Tumor regions (100 slides, listed in **Supplementary Note 1**), Pen mark artifacts (2 slides)
- Pancreas: Islets (1 slide), PanINs (9 slides), Out-of-focus tissue (1 slide)
- Fetal membranes: Red blood cell aggregates (1 slide), Amniotic membrane (1 slide), Chorionic membrane (1 slide)
- Kidney: Normal glomeruli (4 slides), Obsolescent glomeruli (4 slides), Fibrous tissue (4 slides), Tissue artifacts (1 slide)
- Breast: Tissue background (1 slide), Adipose HER2-(1 slide), DCIS with HER2+ (1 slide), Invasive carcinoma HER2+ (1 slide)

These annotations were used for validations. For the interactive classifier training, validations were performed on the same slides as used for the interactive training, unless otherwise specified. Prevalence is defined as the fraction of manually annotated positive pixels within in-tissue tiles.

### Cross validation across donors for age group and anatomical region classification

To compare age groups and anatomical locations, we performed a leave one donor out cross validation procedure. For each iteration, all samples from a single donor were withheld as the validation set. From the remaining donors, samples belonging to one group (either “old” donors or “head of pancreas” anatomical location) were designated as positive, and samples from the comparison group (either “young” donors or “tail of pancreas” anatomical location) were designated as negative for model training. A logistic regression classifier was trained using 500 randomly selected positive patches and 500 randomly selected negative patches from the training set. After training across all leave one sample out folds, a general classifier was constructed by aggregating the coefficients and intercepts from individual models (taking the median across folds) and retaining only the 100 coefficients with the largest absolute magnitude, with all others set to zero. This general classifier was then applied to all positive and negative training samples as well as the held out donor’s samples to obtain predicted probabilities. For each sample, we computed the mean predicted probability, and classification of the left out donor’s samples was determined by comparing these values with the median predicted probabilities for positive and negative training samples, assigning each sample to the group whose median was closest to.

### STQ-imaging pipeline processing

We used version 3.0.0 of the Spatial Transcriptomics Quantification (STQ) [12] pipeline imaging workflow, which features a modular architecture built with Nextflow. This version is optimized for flexibility, scalability, and efficiency, enabling multi-scale imaging feature extraction from H&E- or IHC-stained WSIs.

The pipeline is composed of distinct modules for region of interest (ROI) extraction, image conversion, focus quality control, and standardized output formatting. Its design is extensible, allowing easy integration of new deep learning (DL) models and tools to support evolving spatial biology research. For image analysis, the pipeline incorporates four DL models—InceptionV3 [49], CTransPath [13], MoCoV3 [14], UNI [15], and CONCH [16]—each contributing to a distinct imaging feature set. Unless otherwise specified, we used the CTransPath pretrained model for the analyses in the Results section. Other feature sets based on nuclear morphometry can also be generated with STQ, but were not used in the Results described here.

STQ allows for multi-scale analysis. The pipeline reads image tiles based on a base size defined by the pipeline parameter. These tiles are resized to match the input dimensions required by each DL model using TorchVision’s resize transformation. Users can specify one or more input scales (e.g., 1, 2, 3, 4), which determine tile sizes as multiples of the base. Here, we use scale 1 for all analyses. This allows extraction of features from varying spatial contexts centered on the same coordinate.

Tile overlap is implicitly defined by the base tile size, center-to-center spacing, and grid layout, which can be square, hexagonal, or random. The DIANNE codebase uses center-to-center spacing of STQ as an input for rendering visualizations. Using smaller center-to-center distances results in smoother spatial inference of probability maps.

The five DL models can be used for feature extraction with STQ: InceptionV3, a convolutional neural network pretrained on ImageNet for general image recognition; CTransPath, a transformer-based model trained with contrastive learning on histopathology images; MoCoV3, a self-supervised vision transformer trained using momentum contrast for scalable and efficient representation learning; UNI/UNI2, a self-supervised model trained on over 100 million pathology images to support diverse clinical tasks; and CONCH, a vision-language foundation model trained on image-caption pairs for tasks such as classification, segmentation, and retrieval. All models are implemented in PyTorch, with some (e.g., InceptionV3) also available in TensorFlow. These models were integrated into the STQ pipeline to enable robust and scalable histopathological image analysis.

### MIE pipeline processing

We developed version 1.0.0 of the Multiplex Immunofluorescence Extraction (MIE) pipeline, a modular image processing workflow designed for scalable feature extraction from multiplex immunofluorescence (mIF) whole-slide images (WSIs). MIE is part of the broader Spatial Omics Tools (SOT) framework, which supports automation and reproducibility in spatial omics data analysis.

MIE performs three main tasks: (1) extraction of the region of interest, if requested, (2) tiling of WSIs, (3) tissue detection, (4) extraction of imaging features using a pre-trained foundation model, and (5) unsupervised clustering and visualization for rapid quality assessment. The pipeline is optimized for high-throughput processing and can run on either CPU or GPU, with GPU execution providing a significant speed advantage. The speedup of the feature extraction is at least 15×, depending on the GPU characteristics or number of CPU cores used. We tested 10 CPUs versus NVIDIA Tesla A100 GPU.

For feature extraction, MIE uses KRONOS [17], a vision foundation model trained for spatial omics applications. Tiles are generated from the WSI using a configurable grid layout—square, hexagonal, or random—with user-defined tile size and spacing. Only tiles that contain tissue are processed, as determined by automated tissue detection via OTSU thresholding and OpenCV morphological operations. If a region of interest (ROI) file is provided, only the specified area is cropped with QuPath CLI and analyzed.

After feature extraction, MIE optionally performs a lightweight downstream analysis to provide a quick visual check of data quality. This includes dimensionality reduction using principal component analysis (PCA), followed by construction of a neighborhood graph using correlation distance and 10 nearest neighbors. Clustering is performed at a resolution of 0.35, and the results are visualized in both spatial and UMAP layouts. The downstream analysis parameters along with any analysis parameters of MIE are configurable by the user. The downstream visualizations are not intended for in-depth analysis but serve as a quality check to confirm that the extracted features capture meaningful spatial structure. The primary outputs of MIE are the tiled image grid and the corresponding KRONOS-derived feature matrix, as seen in **Supplementary Fig. 11**, which are intended for user-defined downstream analyses.

### Spatial proteomics data visualization

Multiplex immunofluorescence (mIF) data typically contains more than 3 channels and therefore cannot be directly presented as a 3-channel RGB (red, green, blue) image. DIANNE implements an approach to use the mIF pixel-level values to compute principal components (PC) across all channels and aggregate the PCs into RGB values for display. The PCs can be either left unnormalized, or scaled to normalize the component-wise variances, a choice specified via a DIANNE toolkit parameter. To display all the PCs as RGB values, starting from channel 0 we aggregate every third channel into red channel (R), starting from channel 1 we aggregate every third channel into red channel (G), and starting from channel 2 we aggregate every third channel into red channel (B), clip to low and high quantiles, and normalize the values to a valid range of 8 bit unsigned integer:

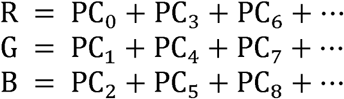

Image transformation is learned within the function “jump-start” and then used in the “annotation” function to ensure consistent visual representation.

### Slide cropping tool

We made a tool for use with STQ to cut out tissue regions before preprocessing, **Supplementary Fig. 12**. It is particularly useful to trim a desired tissue region, remove background, for better display in downstream spatial inference.

## Software and data availability

### Software availability

All software used in this study is publicly available. DIANNE, a software toolkit, documentation and graphical user interfaces, including PCMA (this work), is available at https://github.com/TheJacksonLaboratory/image-differential-annotator. The STQ Imaging Workflow, which provides a workflow for H&E WSI processing including normalization, tiling, and feature extraction [12], can be accessed at https://github.com/TheJacksonLaboratory/STQ. The MIE pipeline of SOT (this work), which performs mIF WSI processing including tiling and feature extraction, is available at https://github.com/TheJacksonLaboratory/spatial-omics-tools.

### Data availability

All datasets used in this study are publicly available. Details of each dataset, including references and descriptions, are provided in Supplementary Table 2. Briefly, the Pediatric Sarcoma H&E WSI dataset [21] comprises over 800 curated and harmonized WSIs of pediatric sarcoma cases. TCGA H&E WSI data were obtained from the National Cancer Institute GDC data portal (https://portal.gdc.cancer.gov/). The CAMELYON16 H&E WSI dataset [23] contains slides of lymph node metastases in breast cancer. The ACROBAT IHC dataset [33], consists of IHC WSIs from female primary breast cancer patients scanned at 10x magnification. Pancreas, kidney, and fetal membranes H&E WSI datasets, as well as pancreas mIF WSI datasets, were obtained from the SenNet Data Sharing Portal (10.60586/SNT254.FDWM.977; 10.60586/SNT645.GFXF.795; 10.60586/SNT389.HBHH.462; 10.60586/SNT536.HKNV.432; 10.60586/SNT753.CKTK.227; 10.60586/SNT922.MBDC.543; 10.60586/SNT336.XGNK.779; 10.60586/SNT227.HLMG.672; 10.60586/SNT484.VLRN.777). Finally, the Pancreas Xenium and matched H&E dataset, comprising 32 datasets of Xenium and matched histology data from the pancreas, is available via 10.60586/SNT793.SZRS.468.

## Acknowledgements

We thank the Single Cell Biology Lab and the Histology and Light Microscopy core of Scientific Services at The Jackson Laboratory. This work was supported by the National Cancer Institute (NCI) PDXNet Data Commons and Coordinating Center U24CA224067 and the National Institute on Aging (NIA) KAPP-Sen Tissue Mapping Collaborative U54AG075941.

The results here are in part based upon imaging data generated by the TCGA Research Network. We acknowledge the use of artificial intelligence services Copilot to revise and edit the text.

These services were not used to generate novel text. Further, we acknowledge the use of GitHub Copilot to edit parts of the code used in this manuscript, which were checked and edited by the authors.

## Author contributions

S.D., J.H.C., J.C.R. and T.S. conceived and designed the study. T.S. annotated whole slide images. S.D. developed software and produced figures and documentation. S.D. and J.H.C. wrote the manuscript. All authors provided insights into the application of the model, contributed to data interpretation, reviewed and revised the manuscript.

