## Supplementary Information for "DIANNE: Segmentation-Free Localization of Histology Differential Attributes"

### Supplementary Figures

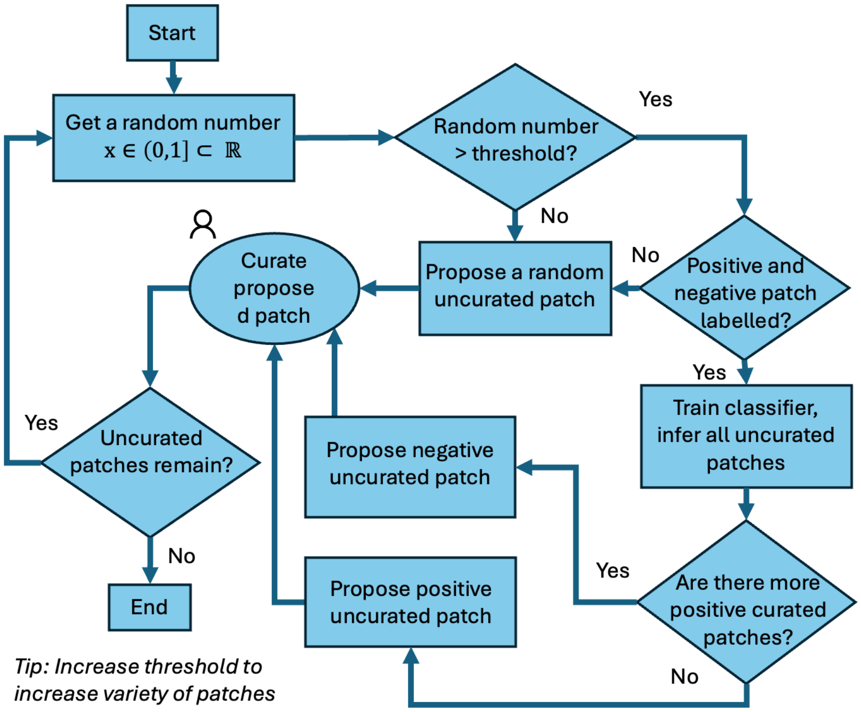


**Supplementary Figure 1. Patch curation proposal algorithm of the Guided GUI.** The active learning algorithm samples a random number x from a uniform distribution on the interval (0,1). If x exceeds a predefined threshold, the system proposes an uncurated patch for classification. Otherwise, a human curator evaluates and labels the proposed patch. Upon successful labeling of both positive and negative examples, a binary classifier is trained to infer labels across all remaining uncurated patches. The system then identifies and prioritizes the most certain samples from each class—specifically, the most likely negative patch and the most likely positive patch—for subsequent human curation. This iterative process continues until no uncurated patches remain. The threshold parameter can be adjusted to modulate the exploration-exploitation balance, with higher thresholds increasing patch diversity at the cost of additional curation effort.

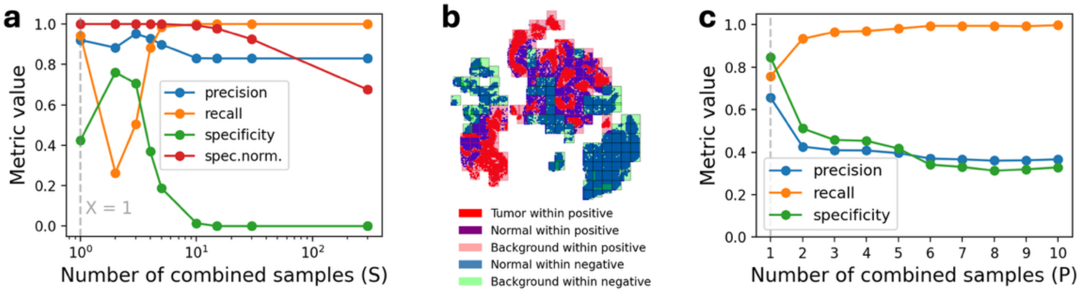


**Supplementary Figure 2. Impact of normal slide or patch aggregation on classification performance.** **a**, Performance metrics as a function of the number of normal slides (S) aggregated to generate a multi-slide normal representation. Specificity declines progressively when S exceeds 3. **b**, Sarcoma slide RMS2341 with colors showing pathologist annotations, at patch resolution. **c**, Performance metrics as a function of the number of normal patches (P) aggregat**ed** to generate a combined normal patch representation**.** Patches are from slide RMS2341. Patch aggregation yields a performance decline when P > 1.

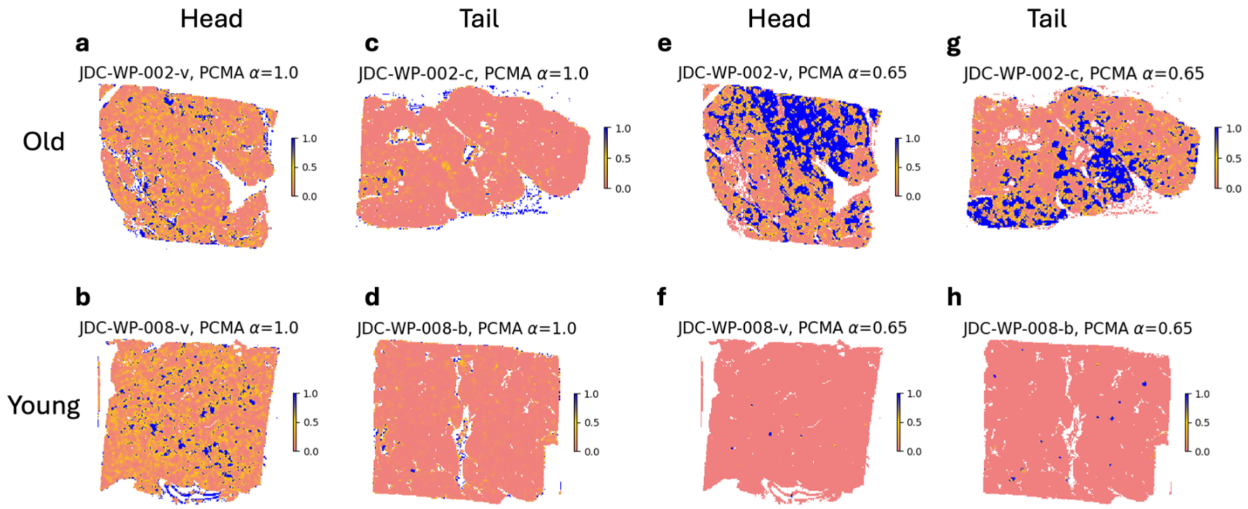


**Supplementary Figure 3. Intraregion heterogeneity for predictors trained on slide-level anatomical region or donor age in pancreas histology images.** Panels **a-d** show predictions of a classifier trained on patches of 17 "pancreas head" tissue slides as positive class, and patches of 15 "pancreas tail" tissue slides as negative class. True status (head/tail, old/young) of the tissue is shown on the axes. Even within head slides, the regions with a high predicted probability of "head" are sparse (dark blue regions, **a** and **b**). Spatial regions with a high probability of "head" within tail slides are also observed (dark blue regions in **c** and **d**), though they are less prevalent than in true head slides. Panels **e-h** show predictions of a classifier trained on patches of 6 "old donor pancreas" tissue slides as positive class, and patches of 10 "young donor pancreas" tissue slides as negative class. Although predicted "old" regions are higher in old slides (**e** and **f**), young slides also contain some regions predicted to be "old" (**g** and **h**).

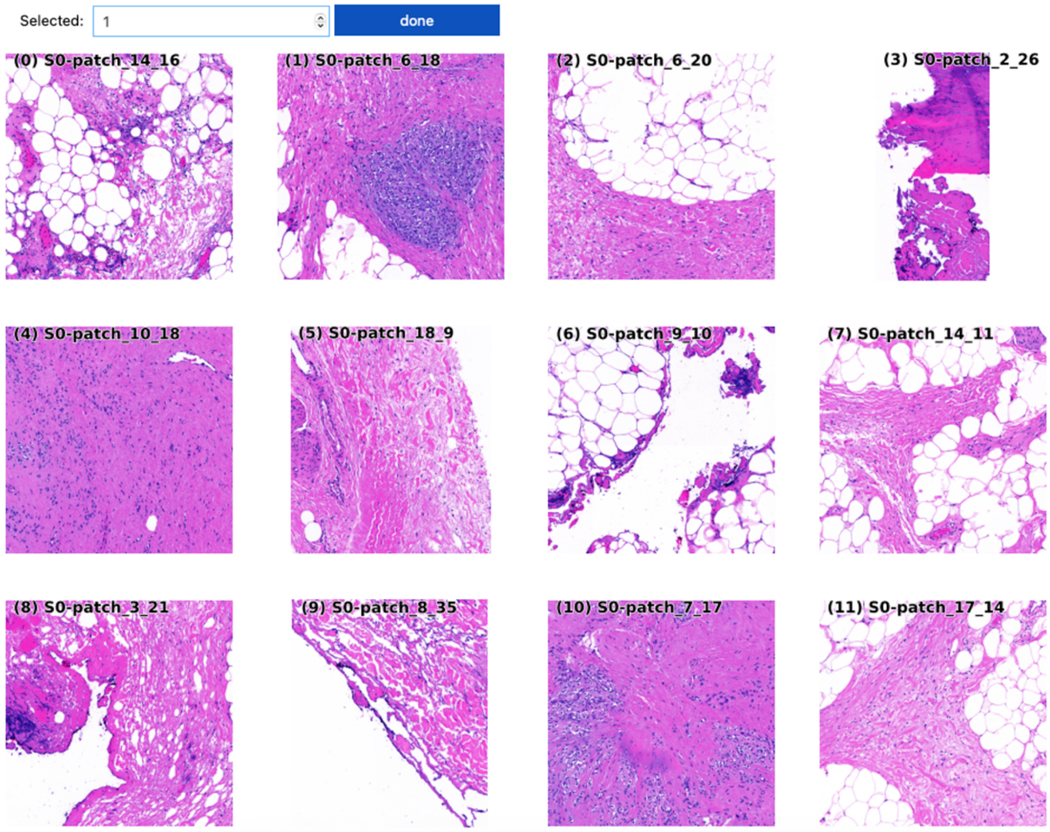


**Supplementary Figure 4. DIANNE Jump-start tool displays patches representing the major classes of imaging patterns in the selected dataset.** This example shows patches from a sarcoma slide**.** After the user specifies the desired numbers of rows and columns, it takes a few seconds to lazy-load the patches into a grid. The operator may enter a number in the box to select a positive patch to begin interactive training.

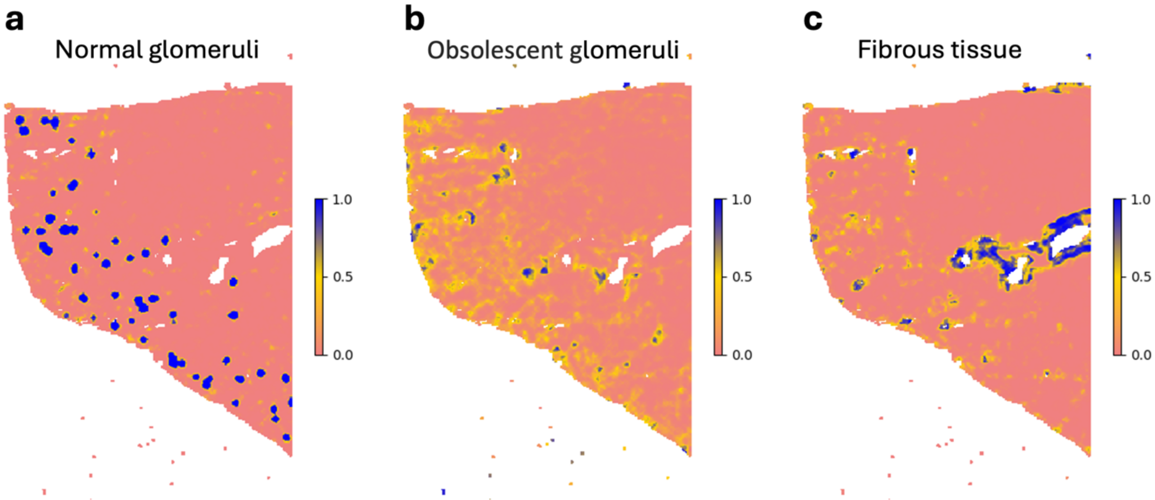


**Supplementary Figure 5. Examples of several regional classifiers trained with DIANNE and applied to a human kidney H&E-stained tissue section from a 49 y.o. male donor. a** Glomeruli, **b** obsolescent glomeruli, **c** fibrous tissues in the renal cortex.

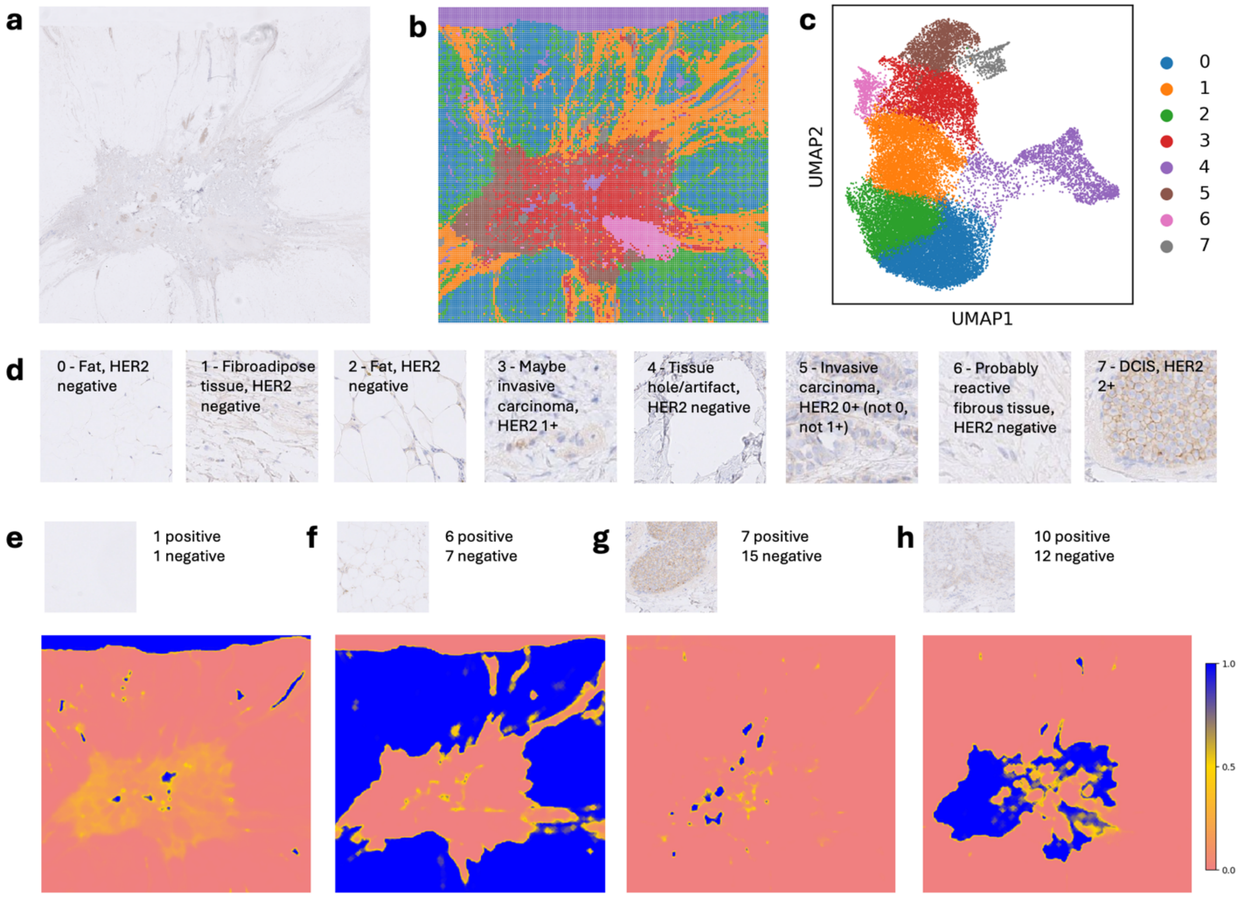


**Supplementary Figure 6. Unsupervised tissue segmentation and spatial search in a HER2-stained breast cancer slide. a**, Whole-slide tissue overview; **b**, spatial map with eight color-coded classes; **c**, UMAP embedding of learned representations with the same eight classes; **d**, representative image patches for each class. **e**–**h** Spatial inference examples for: background, HER2-negative adipose tissue, DCIS with HER2 2+, and invasive carcinoma, respectively. Each example shows the number of positive and negative patches used for training; a positive patch from the training set; and a full-slide probability map where blue indicates high similarity to the reference pattern.

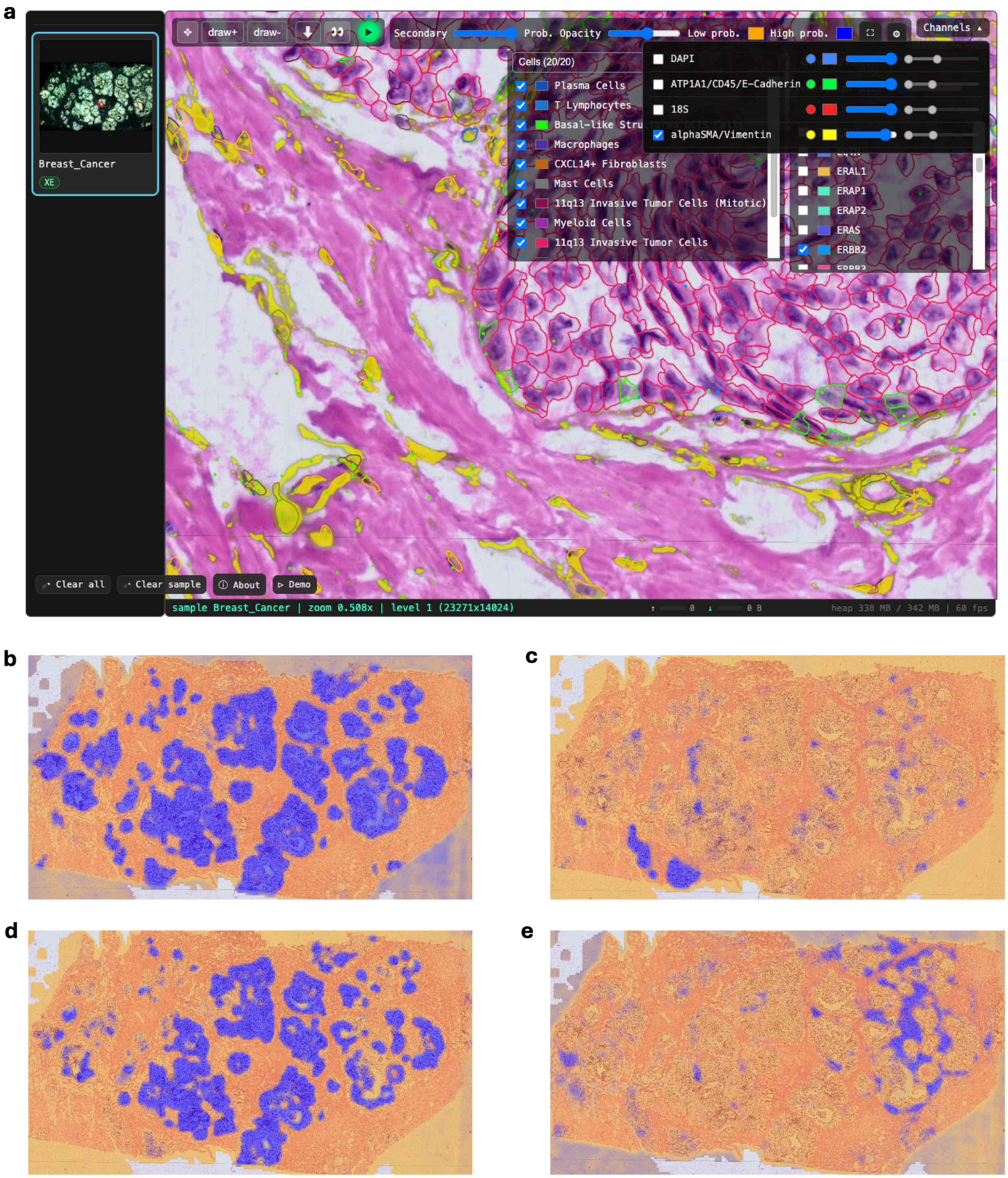


**Supplementary Figure 7. DIANNE interactive analysis of 10x Atera In Situ data. a,** The demo dataset (downloaded from 10x with the cell annotations, H&E image and the Atera bundle) is visualized in the freehand GUI. The mIF stain channel alphaSMA/Vimentin is shown in yellow with opacity 90%. All cells are shown aligned with the H&E and mIF images. Blue and green dots represent ERBB2 and VIM transcripts, respectively. **b,** Tumor classifier trained by freehand annotation of tumor cell-containing regions. In the heatmap, blue indicates high probability and orange low. Classifiers trained to detect basal-like DCIS cells, invasive carcinoma cells, and lymphocyte infiltrated stromal regions are shown in **c-e**, respectively.

**a**

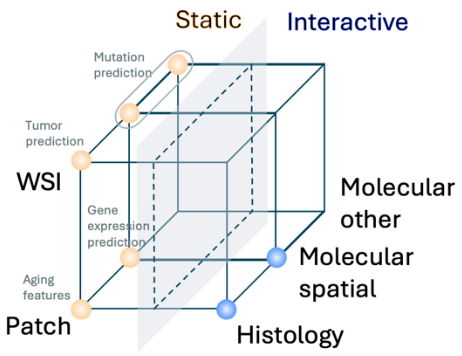


**b**

| Workflow | H&E, mIF, etc. | WSI-based | Patch-based | Other molecular assay | Spatial molecular assay |
|---|---|---|---|---|---|
| Histology static | ✅ | ✅ | ✅ | | |
| Histology interactive | ✅ | | ✅ | | |
| Molecular static | ✅ | ✅ | ✅ | ✅ | ✅ |
| Molecular interactive | ✅ | | ✅ | | ✅ |

**Supplementary Figure 8. Overview of static and interactive workflow variants in DIANNE. a**, DIANNE supports four binary dimensions of analysis (panel a): static versus interactive workflows (left vs. right plane), slide-level versus patch-level resolution (top vs. bottom), and histology versus histology+molecular data modalities (front vs. back). Molecular is further subdivided into spatial (e.g. 10x Xenium In Situ) and non-spatial ("other") categories. **b**, Summary of applicability of workflows. Each workflow can use either H&E, IHC or antibody-based staining.

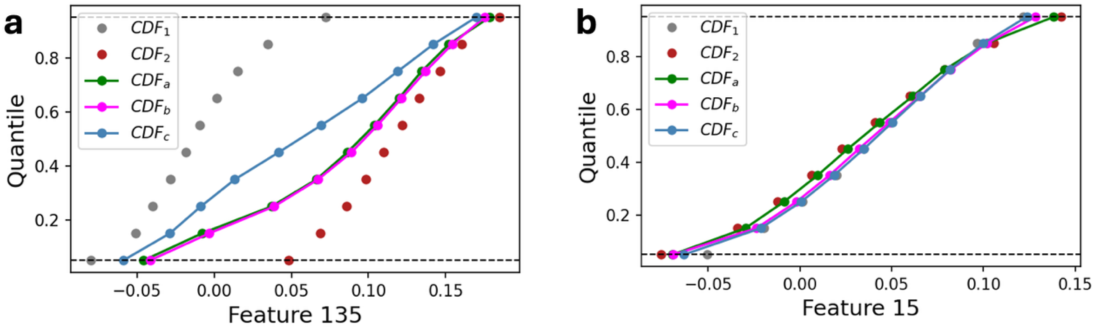


**Supplementary Figure 9. Combining cumulative distribution functions (CDFs) during data augmentation. a**, CDFs for CTransPath feature 135 and **b**, CDF CTransPath feature 15 from combining two samples, normal_005.oid1 and SJDSRCT049192_X1-16699.oid0, which are normal and tumor, respectively. The input CDFs are shown in grey (normal) and red (tumor), while the $CDF_a$ generated from concatenating tile feature values and computing quantiles is shown in green. The combined $CDF_b$ shown in magenta is computed from $CDF_1$ and $CDF_2$ using the weights $\alpha = 0.74$, selected to represent the ratio of tile count in the two samples. By setting $\alpha = 0.5$ we obtain $CDF_c$ shown in blue.

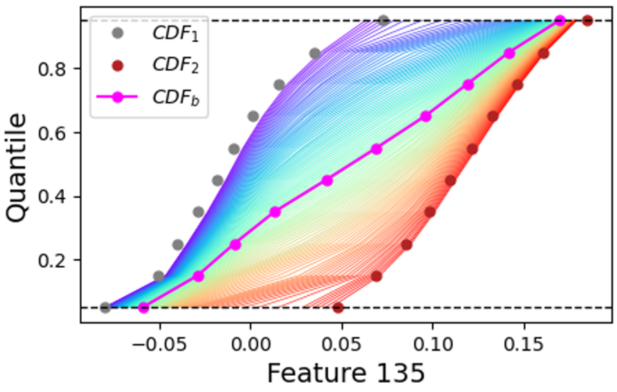


**Supplementary Figure 10. Augmentation of CDFs of CTransPath feature 135 from two samples, normal normal_005.oid1 and tumor SJDSRCT049192_X1-16699.oid0.** The $\alpha = 0.5$ augmented CDF is shown in magenta. Values of $\alpha$ close to 0 correspond to violet and values of $\alpha$ close to 1 correspond to red. These edge cases approach $CDF_1$ and $CDF_2$, respectively.

```
../results/
├── pipeline_info
│   ├── parameters.json
│   ├── execution_timeline_2025-05-05_21-06-04.html
│   ├── execution_report_2025-05-05_21-06-04.html
│   ├── execution_trace_2025-05-05_21-06-04.txt
│   └── pipeline_dag_2025-05-05_21-06-04.svg
├── slide-roi-1
│   ...
└── slide-roi-2
    ├── info.json
    ├── grid
    │   ├── grid.csv
    │   └── grid.json
    ├── features
    │   └── kronos_features.tsv.gz
    ├── figures
    │   ├── spatial_cluster.png
    │   └── umap_cluster.png
    └── downstream-example-data.h5ad
```

**Supplementary Figure 11. MIE output in DIANNE.** The results contain “pipeline_info” with the pipeline parameters and logs, and per-region outputs, with region information, grid files, per-tile features in tabular form, figures, and an AnnData object with clustered data.

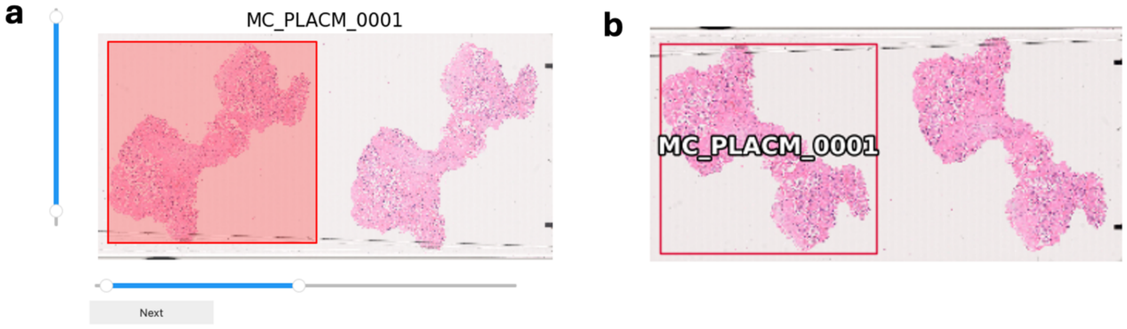


**Supplementary Figure 12. Tissue ROI selection and viewing tools.** The STQ pipeline in the arbitrary grid mode can process a portion of a WSI specified within an input ROI configuration file. See red-shaded rectangle in panel **a**. By default, it spans the entire WSI. The horizontal and vertical sliders in the selection tool, panel **a**, allow users to adjust the ROI selection and instantly see updated selection. The "Next" button advances to the next WSI for ROI selection. Each selection yields a configuration file written in a storage space. To verify the ROI annotations, the viewing tool is used, see panel **b**. The viewing tool can display all selections from all WSIs simultaneously for quick visual inspection.

### Supplementary Tables

**Supplementary Table 1. Slide annotations generated by a pathologist.** For each annotation, the tissue type, staining method, attribute type, sample identifier, and prevalence (proportion of tiles assigned that annotation) are provided. Annotations span multiple tissue types including sarcoma, pancreas, fetal membranes, kidney, and breast, and cover a range of histological and technical attributes such as tumor regions, pen marks, image blur, tissue structures (e.g., islets, glomeruli, amniotic and chorionic membranes), pathological findings (e.g., PanIN, sclerotic glomeruli, DCIS, invasive carcinoma), and technical artifacts.

| *Tissue type* | *Staining* | *Attribute type* | *Sample identifier* | *Prevalence* |
|---|---|---|---|---|
| Sarcoma | H&E | Tumor | Listed below | 76%±21.4% |
| Sarcoma | H&E | Pen marks | PBBMZV.oid0 | 32.0% |
| Sarcoma | H&E | Pen marks | PBBMXT.oid0 | 17.6% |
| Pancreas | H&E | Blurry | JDC-WP-014-aj-HD | 54.5% |
| Pancreas | H&E | Islets | JDC-WP-008-b | 1.5% |
| Pancreas | H&E | PanIN | JDC-WP-008-b | 0.0% |
| Pancreas | H&E | PanIN | JDC-WP-002-v | 0.2% |
| Pancreas | H&E | PanIN | JDC-WP-002-r | 0.4% |
| Pancreas | H&E | PanIN | JDC-WP-012-w | 1.6% |
| Pancreas | H&E | PanIN | JDC-WP-002-ae | 0.5% |
| Pancreas | H&E | PanIN | JDC-WP-008-b-Xe | 0.0% |
| Pancreas | H&E | PanIN | JDC-WP-002-v-Xe | 0.0% |
| Pancreas | H&E | PanIN | JDC-WP-012-w-Xe | 1.8% |
| Pancreas | H&E | PanIN | JDC-WP-002-ae-Xe | 0.3% |
| Membranes | H&E | Red blood cell aggregates | MC-PLACM-0015 | 9.0% |
| Membranes | H&E | Amniotic membrane | MC-PLACM-0015 | 18.9% |
| Membranes | H&E | Chorionic membrane | MC-PLACM-0015 | 20.1% |
| Kidney | H&E | Normal glomeruli | SA-001-L-CM-w | 1.8% |
| Kidney | H&E | Normal glomeruli | SA-001-L-C-a | 6.0% |
| Kidney | H&E | Normal glomeruli | SA-001-L-M-j | 0.0% |
| Kidney | H&E | Normal glomeruli | SA-001-L-P-mnop | 0.0% |
| Kidney | H&E | Sclerotic glomeruli | SA-001-L-CM-w | 0.3% |
| Kidney | H&E | Sclerotic glomeruli | SA-001-L-C-a | 0.1% |
| Kidney | H&E | Sclerotic glomeruli | SA-001-L-M-j | 0.0% |
| Kidney | H&E | Sclerotic glomeruli | SA-001-L-P-mnop | 0.0% |
| Kidney | H&E | Fibrous tissue | SA-001-L-CM-w | 5.4% |
| Kidney | H&E | Fibrous tissue | SA-001-L-C-a | 0.4% |
| Kidney | H&E | Fibrous tissue | SA-001-L-M-j | 3.1% |
| Kidney | H&E | Fibrous tissue | SA-001-L-P-mnop | 38.0% |
| Kidney | H&E | Tissue artifacts | SA-001-L-CM-w | 0.1% |
| Breast | IHC, HER2 | Tissue background | 059-HER2 | 6.0% |
| Breast | IHC, HER2 | Adipose | 059-HER2 | 58.3% |
| Breast | IHC, HER2 | DCIS | 059-HER2 | 0.4% |
| Breast | IHC, HER2 | Invasive carcinoma | 059-HER2 | 13.7% |

**Supplementary Table 2. Data used in this study.** For each dataset, the reference or accession information and a brief description are provided, including the imaging modality, tissue type, and, where applicable, patient demographic information.

| **Dataset Name** | **Reference** | **Description** |
|---|---|---|
| Pediatric Sarcoma H&E WSI | 10.1101/2025.06.10.25328700 | Over 800 curated and harmonized WSIs of pediatric sarcoma cases |
| TCGA H&E WSI | https://portal.gdc.cancer.gov/ | National Cancer Institute GDC data portal |
| CAMELYON16 H&E WSI | 10.1001/jama.2017.14585 | Diagnostic Assessment of Deep Learning Algorithms for Detection of Lymph Node Metastases in Women with Breast Cancer |
| ACROBAT, IHC | 10.1016/j.media.2024.103257, https://researchdata.se/en/catalogue/dataset/2022-190-1 | IHC WSI from female primary breast cancer patients scanned at 10x magnification |
| Pancreas H&E WSI | 10.60586/SNT254.FDWM.977 | Histology data from the pancreas of a 23-year-old female, JDC-WP-008-b |
| Pancreas H&E WSI | 10.60586/SNT645.GFXF.795 | Histology data from the pancreas of a 69-year-old white female, JDC-WP-012-ae |
| Pancreas H&E WSI | 10.60586/SNT389.HBHH.462 | Histology data from the pancreas of a 69-year-old white female, JDC-WP-012-w |
| Pancreas H&E WSI | 10.60586/SNT536.HKNV.432 | Histology data from the pancreas of a 71-year-old white female, JDC-WP-002-r |
| Pancreas H&E WSI | 10.60586/SNT753.CKTK.227 | Histology data from the pancreas of a 71-year-old white female, JDC-WP-002-v |
| Kidney H&E WSI | 10.60586/SNT922.MBDC.543 | Histology data from the kidney (left) of a 49-year-old white male, SA-WLK-019-w |
| Placenta H&E WSI | 10.60586/SNT336.XGNK.779 | Histology data from the placenta of a 34-year-old asian female, MY-PW-0015-PLACM-FO |
| Pancreas mIF WSI | 10.60586/SNT227.HLMG.672 | PhenoCycler data from the pancreas of a 69-year-old white female, JDC-WP-012-ae |
| Pancreas mIF WSI | 10.60586/SNT484.VLRN.777 | PhenoCycler data from the pancreas of a 69-year-old white female, JDC-WP-012-w |
| Pancreas Xenium and H&E | 10.60586/SNT793.SZRS.468 | 32 datasets of Xenium and matched histology data from the pancreas |
| Breast cancer Atera In Situ | 10x Genomics | 10x Genomics demo dataset |

### Supplementary Notes

**Supplementary Note 1. Sarcoma 100 annotated slide identifiers**: 0b6d0168-d599-4f97-b096-543abd938b3c.oid2, 0d2deb29-55c8-4b78-a183-98213ab734a2.oid0, 0d2deb29-55c8-4b78-a183-98213ab734a2.oid3, 16088.oid0, 17015.oid0, 19300.oid1, 20028.oid0, 20076.oid0, 224dda13-692f-4003-9d4e-e841649ed647.oid1, 23013.oid0, 3f81389c-1962-4821-a057-f81ff625a193.oid0, 455a896d-fa56-4a89-9601-b3a7c84e0786.oid1, 592ea68a-cb88-422f-bae9-a010f7d663b2.oid2, 689722ac-f2ac-4a5f-9453-32a2cb321a87.oid0, 6b4a6d67-4aa2-40b3-a1ad-72c8101baddd.oid1, 88a11481-3071-4f7f-b6af-a969d0248ff6.oid0, PBBLCW.oid0, PBBMXY.oid0, PBBNBW.oid1, PBBNDK.oid0, PBBPDU.oid1, PBBVHA.oid0, PBBYFL.oid0, PBBYIV.oid1, PBBYKH.oid0, PBBZAE.oid0, PBBZLW.oid0, PBCAEG.oid1, PBCAKP.oid0, PBCBKT.oid0, PBCBMD.oid0, PBCCCI.oid0, RMS2134.oid0, RMS2140.oid1, RMS2148.oid0, RMS2172.oid0, RMS2173.oid0, RMS2187.oid0, RMS2194.oid0, RMS2198.oid0, RMS2203.oid0, RMS2234.oid1, RMS2248.oid0, RMS2269.oid0, RMS2280.oid0, RMS2291.oid0, RMS2296.oid0, RMS2308.oid1, RMS2310.oid2, RMS2323.oid0, RMS2328.oid0, RMS2328.oid1, RMS2330.oid0, RMS2332.oid0, RMS2338.oid0, RMS2339.oid0, RMS2341.oid0, RMS2345.oid1, RMS2350.oid0, RMS2367.oid0, RMS2379.oid0, RMS2396.oid0, RMS2423.oid0, RMS2444.oid0, RMS2445.oid0, RMS2448.oid0, RMS2451.oid0, RMS2453.oid0, RMS2458.oid0, RMS2466.oid0, RMS2476.oid0, RMS2477.oid0, RMS2482.oid0, RMS2486.oid0, RMS2507.oid0, RMS2507.oid1, SJDSRCT046155_X1-16397.oid0, SJHGS030456_R1-7485.oid0, SJIFS030375_X1-16379.oid1, SJRHB010468_X1-16300.oid0, SJRHB010928_X1-16425.oid0, SJRHB012405_X1-16322.oid0, SJRHB012405_X1-16322.oid1, SJRHB012_D-6811.oid1, SJRHB012_X-16418.oid0, SJRHB013759_X1-16323.oid0, SJRHB030549_X1-16434.oid0, SJRHB030550_X1-16368.oid2, SJRHB030765_X1-16430.oid1, SJRHB030787_X1-16429.oid0, SJRHB031320_X1-16426.oid1, SJRHB031691_D2-6978.oid0, SJRHB031691_X3-16309.oid1, SJRHB049188_D1-6809.oid0, SJRHB071779_X1-16498.oid1, SJSS063828_X1-16345.oid0, SJSTS030383_X1-16313.oid0, SJSTS031700_D1-6832.oid0, a674f8cc-8e0f-448b-a061-93ca90ea92eb.oid0, da3410a7-8ff0-4d28-9454-21006d0af689.oid1
